# Cross-reactive TCR with alloreactivity for immunodominant HIV-1 epitope Gag TL9 with enhanced control of viral infection

**DOI:** 10.1101/2021.06.06.447276

**Authors:** Yang Liu, Dan San, Lei Yin

## Abstract

Although both HLA B*81:01 and HLA B*42:01 are members of the B7 supertype and can present many of the same HIV-1 epitopes, the identification of a dual-reactive T-cell phenotype was unexpected, since structural data suggested that TL9 peptide binds to each allele in a distinct conformation. How the dual-reactive TCR recognizes these radically distinct p-MHC surfaces is revealed by our structural study, that the introduction of TCR T18A induces a molecular switch of the TL9 peptide in B4201 to approach its conformation in B8101. Most importantly, unique docking of CDR3β towards MHC but not peptide ligand strengthens the peptide tolerance of T18A, extends the ability of TCR to adapt mutations. Moreover, the high affinity of dual-reactive TCR for WT and escape mutant TL9 highlights the functional advantage of the alloreactive phenotype.

## INTRODUCTION

Antigen-specific T cell immunity is a fundamental ‘law’ of immunology, that is, T cell responses are highly specific and are developmentally restricted to the recognition of self-HLA(Jameson, S.C., Hogquist, K.A., and Bevan, 1995; R M Zinkernagel, 1974a) by T cell receptor (TCR). Most T cells recognize only certain antigens presented by certain host-derived HLA molecules. However, as we previously described, T cells are often cross-reactive with different antigens and different HLAs. Some T cells can break the restriction of HLA and can also react directly with HLA molecules from unrelated individuals(Colf et al., 2007; Felix and Allen, 2007; L A Sherman, 1993), which is called ‘alloreactivity’ and can induce extra immune responses. Such alloreactivity is harmful to transplanted cells that patients with some HLA mismatches can have severe T cell immune responses and result in poor results of transplantation, known as taboo mismatches(Doxiadis et al., 1996; Kawase et al., 2007). And many pieces of evidence showed T cell cross-restriction is a major cause of tissue transplant-related morbidity and mortality(K Fleischhauer, N A Kernan, R J O’Reilly, B Dupont, 1990; Macdonald et al., 2003; Mifsud et al., 2008).

How T cell receptor recognizes MHC and peptide and how they play the vital roles in controlling diseases or inducing diseases attracts popular interests(R M Zinkernagel, 1974b)^−13^. As for alloreactivity reactions, most of the researches aims at the injury they induced for self-tissues. We wondering if alloreactivity reactions could play good roles naturally or even artificially. Similar situations have been noted by our previous studies in other kinds of cross-reactivity. We previously reported that the T cell with the same TCR could be cross-reactive to both MHC I and MHC II positive cells. In some HIV patients CD8^+^ T cells that are trained to recognize MHC I with their TCR were turned to recognize MHC II since CD4^+^ T cells had been hugely destroyed by the virus. Also in the case of tumor immunotherapy, tumor-reactive T cells can be cross-reactive with altered tumor antigen. And when cross-activated these T cells can kill tumor cells and be protective from the tumor. Recently, A subset of T cells that cross-recognized the TL9 epitope bound by B*81:01 or B*42:01 alleles was identified in HIV-infected people(Ogunshola et al., 2018), despite the absence of one allele. And these cross-reactive T cells are correlated with the better outcome for HIV-infected patients, which showed the potential for clinical therapy. Why this alloreactivity happened and how it can be protective from HIV attracts our interests.

Although multiple HLA-B alleles can present the TL9 epitope, the frequency and pattern of TL9 epitope mutations are distinct, and have different effects on HIV-1 replication ability(Edwards et al., 2002; Frater et al., 2007; Leslie et al., 2006a; Ntale et al., 2012). Several explanations were raised for the differential selection pressure exerted on HIV-1 by closely related HLA alleles, including various TCR clonotype usage, different TCR affinities resulting in different cross-recognition properties for TL9 variants(Geldmacher et al., 2009a; Kløverpris et al., 2016; Leslie et al., 2006b), and the completely distinct interact surface presented by TL9 in HLA-B*81:01 and HLA-B*42:01(Kløverpris et al., 2015a). A phenomenon is suggested by these factors: there are different escape pathways of HIV-1 to adapt to different selection pressures when confronted with the CD8^+^T cell response targeting the same epitope but restricted by different HLA molecules. At a population level, this may result in differential HLA-associated viral replication capacity and disease prognosis(Carlson et al., 2012).

In this study, we investigate the mechanism of the high-affinity CD8^+^T cell response to immunodominant HIV-1 epitope Gag-TL9 by first reporting its TCR-pHLA ternary-complex structure. In addition, the cross-restriction structure of the same TCR was determined, showing the T18A adopts very similar binding orientations although the conformation of the peptide Gag-TL9 are largely different when Gag-TL9 bound to its host-selecting B*81:01-TL9 and allogeneic B*42:01-TL9 molecules. To be cross-reactive with both, CDR3β of T18A adapts a rare docking position over the conservative MHC surface to avoid contacting the peptide. Moreover, an unusual open form of Vα (the β sheet usually formed by Jβ and Vβ are not formed) was used for recognizing both alleles, and interestingly this unusual open form was firstly reported for cross-reactively interacting with MHCI and MHC II in our previous study(Yin et al., 2011). In the context of this unusual alloreactive TCR, TL9 peptide exhibited dramatic plasticity when bound to B*42:01 upon TCR introduction and adopt a new conformation closer to its configuration in B*81:01 context, which indicates an induced-fit molecular adaptation mechanism for recognizing conformational different peptides. Thus the ability of the high tolerance for recognizing altered peptides might ensure the recognition for the mutants of TL9 and maintain the immune responses for HIV. Therefore, the study of this naturally unexpected cross-reactivity highlights the fundamental basis for alloreactivity in chronic virus infection and key points for controlling HIV by the immune system.

## RESULT

### Overview of the crystal structure of T18A/HLA-B*81:01/TPQDLNTML and T18A/HLA-B*42:01/TPQDLNTML complex

To critically examine why T18A TCR can creatively bind to distinct antigen-presenting surfaces in different HLA contexts, we determined the structure of T18A and TPQDLNTML in the B*81:01 and B*42:01 complexes. The statistics of the crystals were described in table S1, and the structures of the ternary complexes were shown (Figure 1A, C). The T18A TCR combines pMHC in a traditional diagonal manner, with a total buried surface area(Lesk and Chothia, 1987) (BSA) of 1732.6 and 1613.1 Å^2^ in B*81:01 and B*42:01 background, respectively, which fell within the range of known BSA(Rossjohn et al., 2015). The relative contact footprints of the complementarity determining region (CDR) loops at the TCR-pHLA interfaces were very similar (Fig.1B, D). Not all of the CDR loops contribute equally to the interaction, CDR3α, CDR3β, CDR2β dominate the interaction on the TCR-pMHC interface of both complexes. In the T18A/HLA-B*81:01/TPQDLNTML complex, the dominant contribution of Vβ domains was observed at the interface (Vα 40.2%; Vβ 59.7%). In the T18A/B*42:01/TPQDLNTML complex, different involvement of variable domains (Vα 38.4%; Vβ 61.5%) was observed. There was one salt bridge (CDR3β-D100 with B*81:01-R153, CDR3β-D100 with B*42:01 R153) at the interface of both complexes, but the T18A/B*81:01/TPQDLNTML complex formed more hydrogen bonds (Table S2). TL9 peptides contribute 16% to the BSA in the B*81:01 complex, and 15 % in the B*42:01 complex.

**Figure 1.**
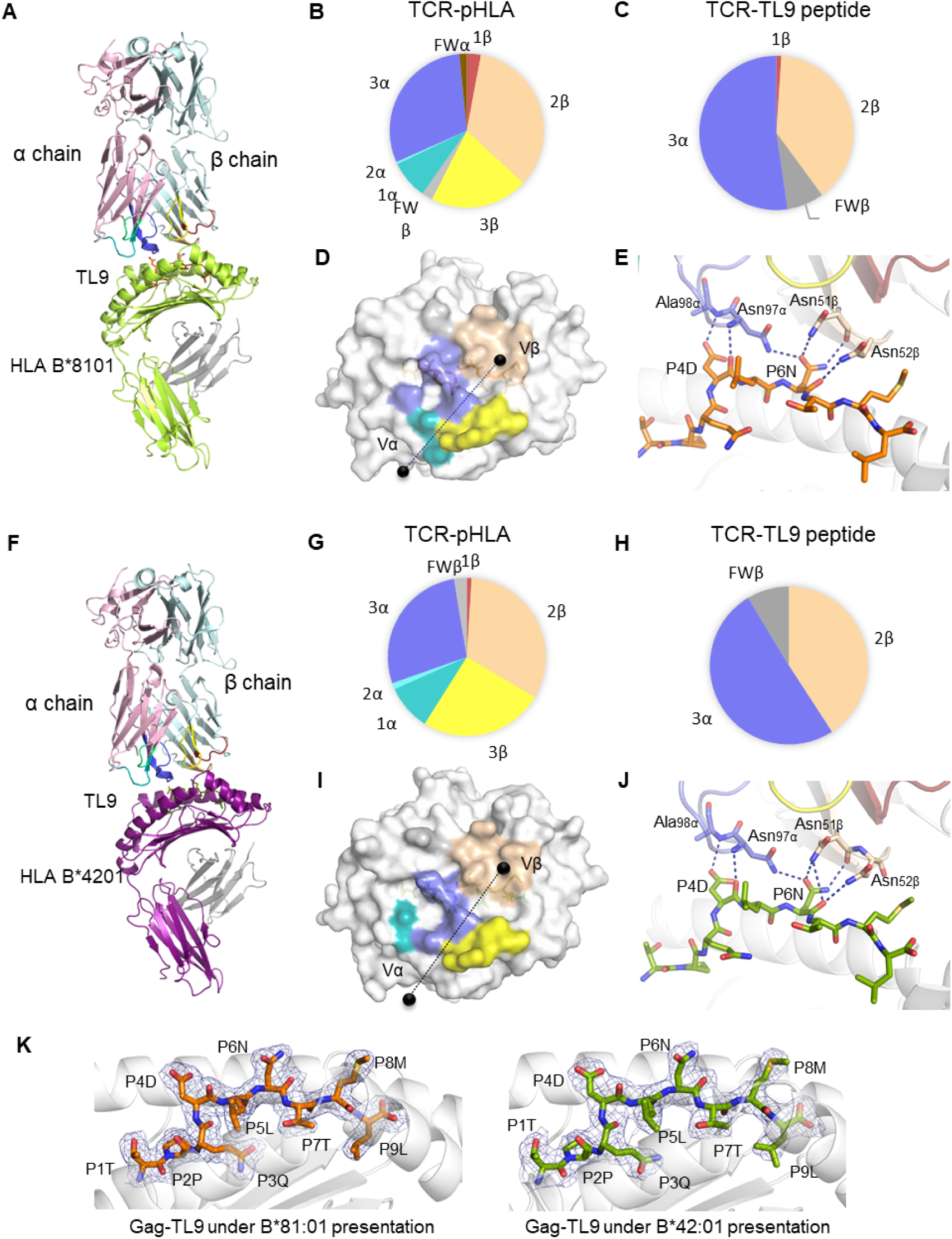
T18A TCR cross-recognition of the TL9 epitope presented by HLA-B8101 and HLA-B4201 alleles. (A). The T18A TCR (T18Aα in pale pink, T18Aβ in pale cyan) recognize TL9 presented by HLA-B8101. Heavy chain of HLA-B8101, and HLA-B4201 are shown in limon, and purple, respectively. The CDR1α, CDR2α, and CDR3α loops are shown in teal, limegreen, and blue, whereas the CDR1β, CDR2β, and CDR3β loops are shown in firebrick, light orange, and yellow, respectively. (B) Pie charts show the contribution of TCR segments toward the pHLA complex. (C). Interactions of TCR towards peptide. (D). The footprint of T18A TCR on the surface of HLA-B8101-TL9 complex. The colors correspond to TCR segment showed in pie chat; the center mass of Vα and Vβ domains were represented by black spheres. (E). Detailed interactions of T18A TCR with Gag-TL9 epitope in the context of HLA-B8101. Blue dashes denote hydrogen bonds; peptide amino acids are indicated in single-letter abbreviations and TCR residues are labeled in three-letter abbreviations. The colors correspond to TCR segment showed in pie chat. (F) The T18A TCR recognize TL9 presented by HLA-B4201. (G-J) Similar profiles as (B-E) of T18A TCR but on the surface of HLA-B4201-TL9. (K). Refined maps (2Fo-Fc) of the peptide in HLA-B complexes. The HLA molecules are represented in cartoon, and the peptides are represented as stick. The online version of this article includes the following source data for figure 1: **Table S1.** Data collection and refinement statistics of TCR-peptide-HLA complexes. **Table S2.** Contact table of T18A/HLA-B*81:01/TL9 and T18A/HLA-B*42:01/TL9

The CDR3α and CDR2β of the T18A sit above the peptide in both complexes and dominate the interaction between TCR and peptide (CDR3α 52%, CDR2β 39% in B8101, CDR3α 50%, CDR2β 41% in B4201). T18A adapts a docking angle of 43° across the antigen-binding groove in both complexes, and no dramatic sliding of the TCR on the pMHC surface is found. To conclude, although the conformation of TL9 differs under the restriction of B81 or B42, and the polymorphism of B81 and B42 α-helical residues also contributes to the different contact surfaces, TCR T18As adopt similar positions at the pMHC interface across B*81:01 and B*42:01 restriction.

In the interaction between T18A TCR and HIV-1 Gag-TL9 epitope, CDR2β and CDR3α loops were the main contributors, which was characterized by strong hydrophilic interactions involving multiple asparagine. In the T18A/B*81:01/TPQDLNTML complex (Fig.1E), CDR2β and CDR3α loop dominated TCR-peptide interactions, and formed six hydrogen bonds with the peptide. Among them, the Asn97 and Ala98 residues of CDR3α loop formed a hydrogen bond network with the P4D of the TL9 peptide, while Asn97 formed a hydrogen bond with the side chain of P6N. The Asn51 and Asn52 of CDR2β loop form three hydrogen bonds with the side chain and backbone of P6D. In the T18A/B*42:01/TPQDLNTML complex (Fig. 1J), the configuration of the peptide was almost the same as that of under B*81:01 presentation, which results in similar hydrophilic TCR-peptide-interactions. To further confirm this interesting observation, the electron density maps of the TL9 in B8101 and B4201 presentation upon TCR binding were shown and compared (Fig 1K).

### ‘Induced-fit’ mechanism of the TL9 peptide presentation upon TCR binding

Next, we aimed to observe the configuration change of TL9 peptide before and after TCR accommodation. HIV Gag-TL9 epitope exhibits distinct conformations when presented by B8101 versus B4201 (Fig. 2B), but adapts similar conformation after TCR binding in the context B8101 and B4201 based by our structural evidence (Fig. 2C).

**Figure 2.**
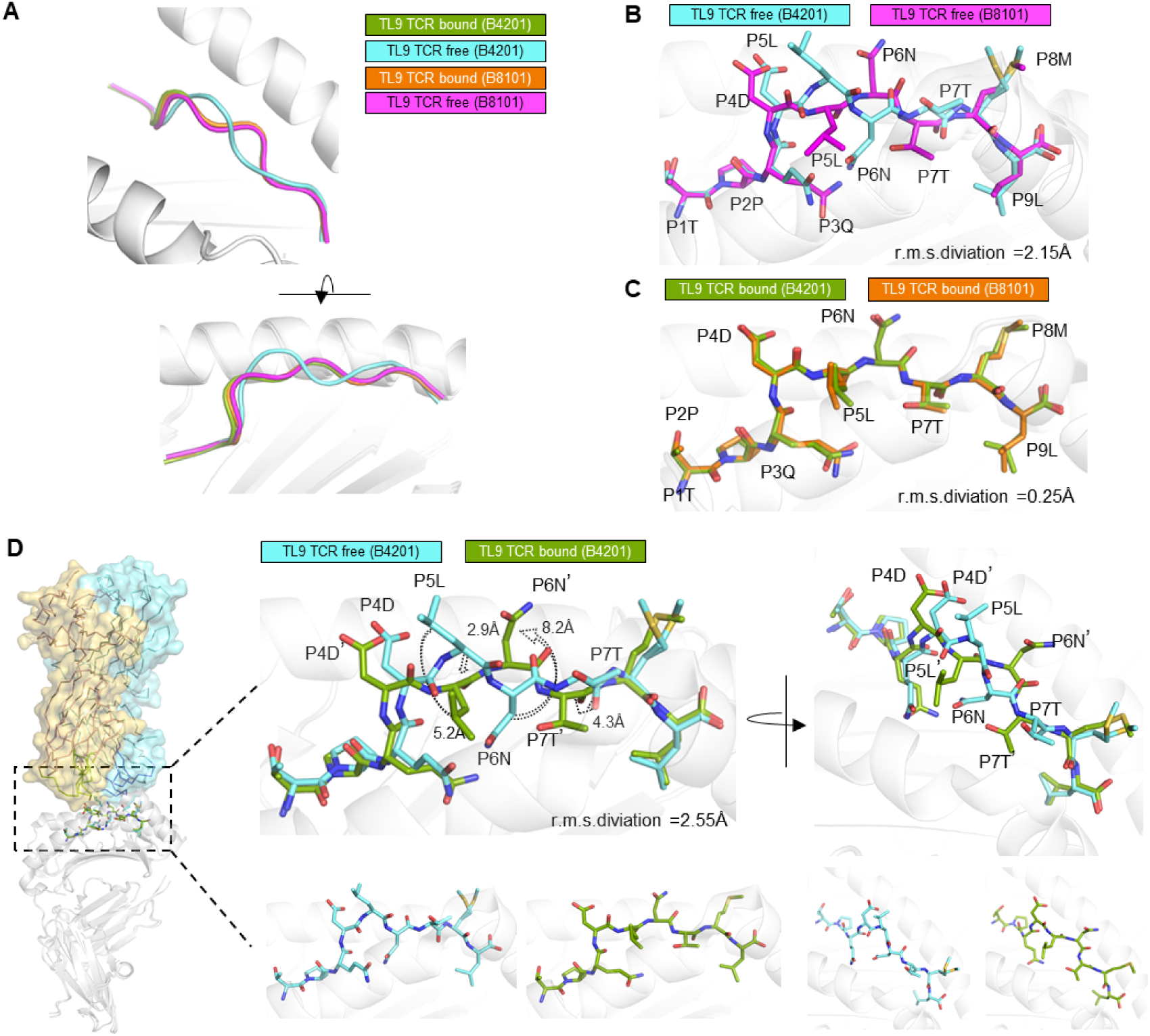
In the context of B4201, T18A recognize the TPQDLNTML peptide very distinctly by inducing a shifting of peptide register and return the TL9 peptide to its B8101 conformation. (A) The register change of TL9 peptide seems due to TCR binding make its conformation closer to its B8101 register. (B) HIV Gag-TL9 epitope exhibits distinct conformations when presented by B8101 versus B4201 (PDB: 4U1I,4U1J(Kløverpris et al., 2015a)). (C) TL9 peptide adapts similar conformation after TCR binding when presented by B8101 and B4201. (D) The diagram of TCR-binding-induced TL9 register shift in the B4201 restriction. The side chain and backbone of P5L is pressed into the bind groove for about 5 Å, the solvent exposed P7T is also press to the bind groove for 4.3 Å. The buried residue P6N shifts upwards by 8.2 Å and contact to the CDR2β of T18A. The online version of this article includes the following source data for figure 2: **Figure S1.** Comparison of free or TCR-bound TL9 peptide when presented by HLA-B*81:01.

Under the B*42:01 restriction, the electron density showed that the central part of TPQDLNTML had a ‘conformational switch’ compared to its conformation in the free pMHC (Fig. 2D). The side chain of leucine at P5 (P5L) turned down with a movement of about 5.2 Å, and its peptide backbone was pressed toward the antigen-binding cleft. At the same time, anchor residue P6N was flipping by 112°, becoming solvent exposed and was involved in CDR2β interactions. On the contrary, solvent exposed P7T shifted towards the base of the groove by 4.3 Å and acted as a secondary anchor residue. The ‘molecular-switch’ resulted in a less bulged conformation of TL9 peptide. From a structural perspective, TCR forced the side chain of the most exposed P5L and backbone of P4-P5 to embed towards the antigen-binding cleft, and popped out the asparagine up and out of the binding groove. Remarkably change in peptide configuration was reflected at r.m.s. deviation of 2.55 Å when the bound and free HLA-B*42:01 TL9-binding domains were superimposed.

In contrast, the configuration of free TPQDLNTML and TCR-bound TPQDLNTML was almost identical under the B81 restriction (Fig.S1). Collectively, in the B42 context, TCR binding induced ‘conformational switch’ and made the backbone configuration much closer to, the conformation of TL9 peptide under B81 restriction (Fig. 2A). The structural rearrangement of peptide occurred upon T18A binding resulted in closer p-MHC surface and similar adaptation of the TCR across B42 and B81 restriction, thus an ‘induced fit molecular mimicry’(Macdonald et al., 2009) might underpinned this cross-restriction structurally.

It was intriguing to investigate how the prominent bulge of TL9 peptide in the free pMHC of the B42 context, was pressed into the antigen-binding groove upon TCR engagement. The superimpose of unbound and bound TL9/HLA-B4201 complexes (Fig. 3A) confirmed that the clashes with CDR3α loop drive the conformational switch of the peptide. Clashes on peptide involved the side chain of P4D and both backbone and side chain of P5L, which competed with Asn96 and Asn97 of CDR3α of TCR. The most serious clashing occurred between the side chain of P5L and the side chain of Asn97, which both occupied the same volume. As TCR and peptide ligands both owned certain extent of plasticity, we wonder why it was peptide itself to adapt to TCR accommodation but not in reverse. A net of hydrogen bonds was observed in the bottom end of CDR3α loop, which fixed the structure of CDR3α backbone. Moreover, W149, K148, T145 and Y86 of HLA-B4201 formed a salt bridge and three hydrogen bonds with P9L and P8M at the TL9 C-terminus. The strong anchoring of C-terminal of the peptide limited the conformational switch in the middle of the peptide, rather than an extending of the peptide C-terminus. To conclude, both relative rigid CDR3α loop and C-terminal anchoring induced the conformational switch of TL9 after T18A involving.

**Figure 3.**
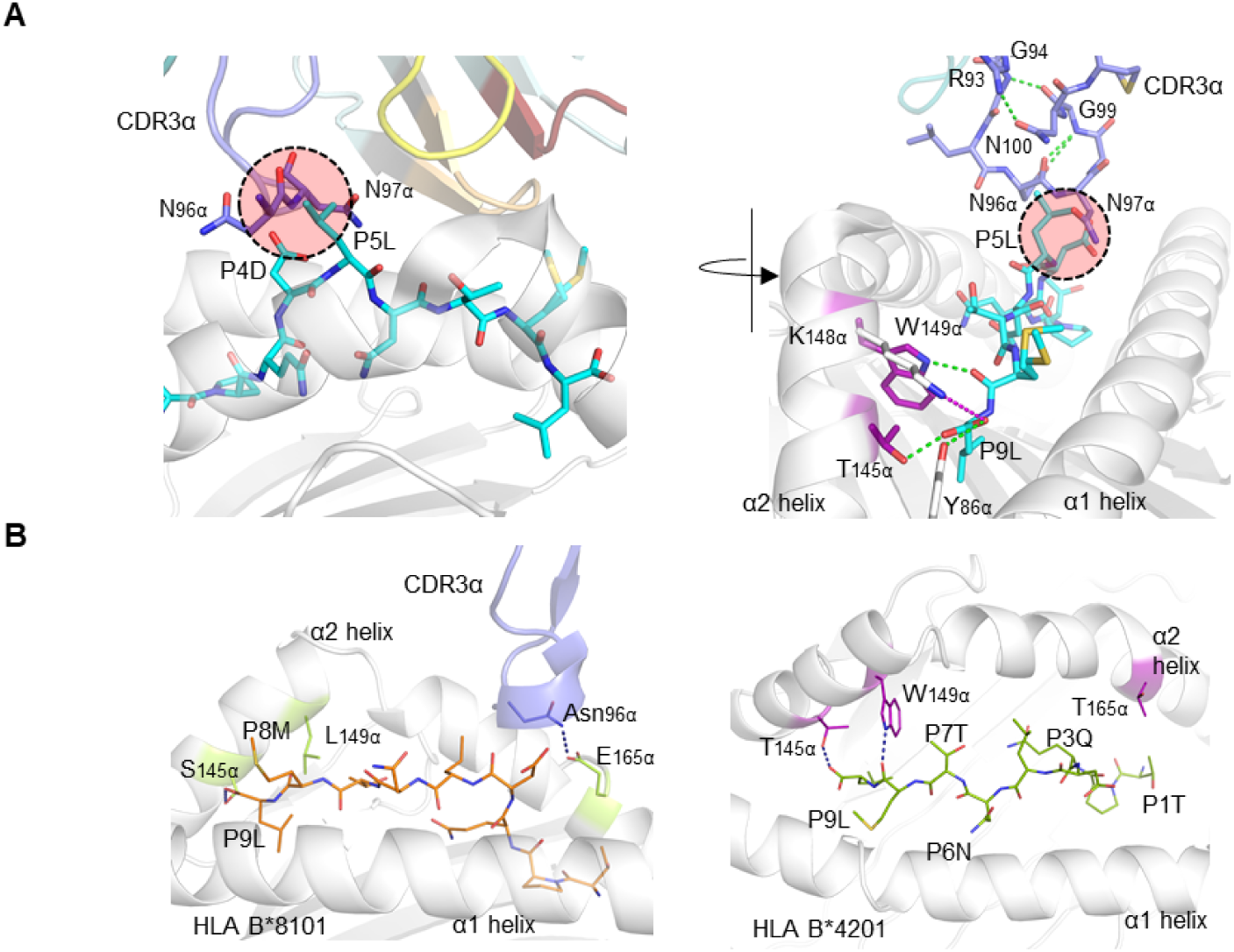
Explanation of the induced-fit mechanism in TL9-B4201 recognition by T18A TCR. (A) Left panel: Steric clashes between peptide (cyan) N-terminal P4D, P5L and Asn96, Asn97 of CDR3α (blue). Right panel: hydrogen bonds matrix increases CDR3α rigidity and drives the peptide to adapt TCR accommodation, instead of TCR to adapt to peptide. And the anchoring of peptide C terminus by W149α, K148α, T145α, and Y86α also contribute to conformational change in the TPQDLNTML peptide upon TCR involving. (B) Illustration of the HLA-B polymorphism in the peptide binding groove. Compare to S145α, L149α in B8101, T145α, W149α in B4201 anchor the C terminus of peptide tighter than B8101 and contribute to conformation change of peptide register. The green dashed lines represent hydrogen bonds and the purple dashed lines represent salt bridges.

In addition, we also compared the effect of HLA polymorphism on TCR binding in the T18A TCR system. HLA-B*81:01 and HLA-B*42:01 are two popular alleles in the African population, differ by 5 residues, of which 3 are located in the peptide-binding groove and may contribute on the interaction (Fig. 3B). Compared with L149 and S145 of HLA-B*81:01, W149 and T145 of HLA-B*42:01 had stronger interactions with TL9 peptide, which fixed the peptide C-terminus and contributed to the conformational adaptation of peptide upon T18A binding. However, E165 of HLA-B*81:01 formed hydrogen bonds with Asn96 of T18A CDR3α, contributed to a stronger TCR-MHC interaction than HLA-B*42:01.

### A slightly twisting of T18A CDR loops enables adaptation to polymorphic MHC ligands

Structures of T18A TCR with B8101-TL9 or B4201-TL9 were superimposed by MHC, although the configuration of TCR was almost similar, a small pivoting about 1 Å was observed mainly on CDRα loops but not CDRβ loops (Fig. 4A-C). To investigate which factor will cause this twisting, CDRα residues and adjacent atoms were overlaid (Fig. 4D), and the polymorphic residue E165 of B8101 and T165 of B4201 was coincidental in this region.

**Figure 4.**
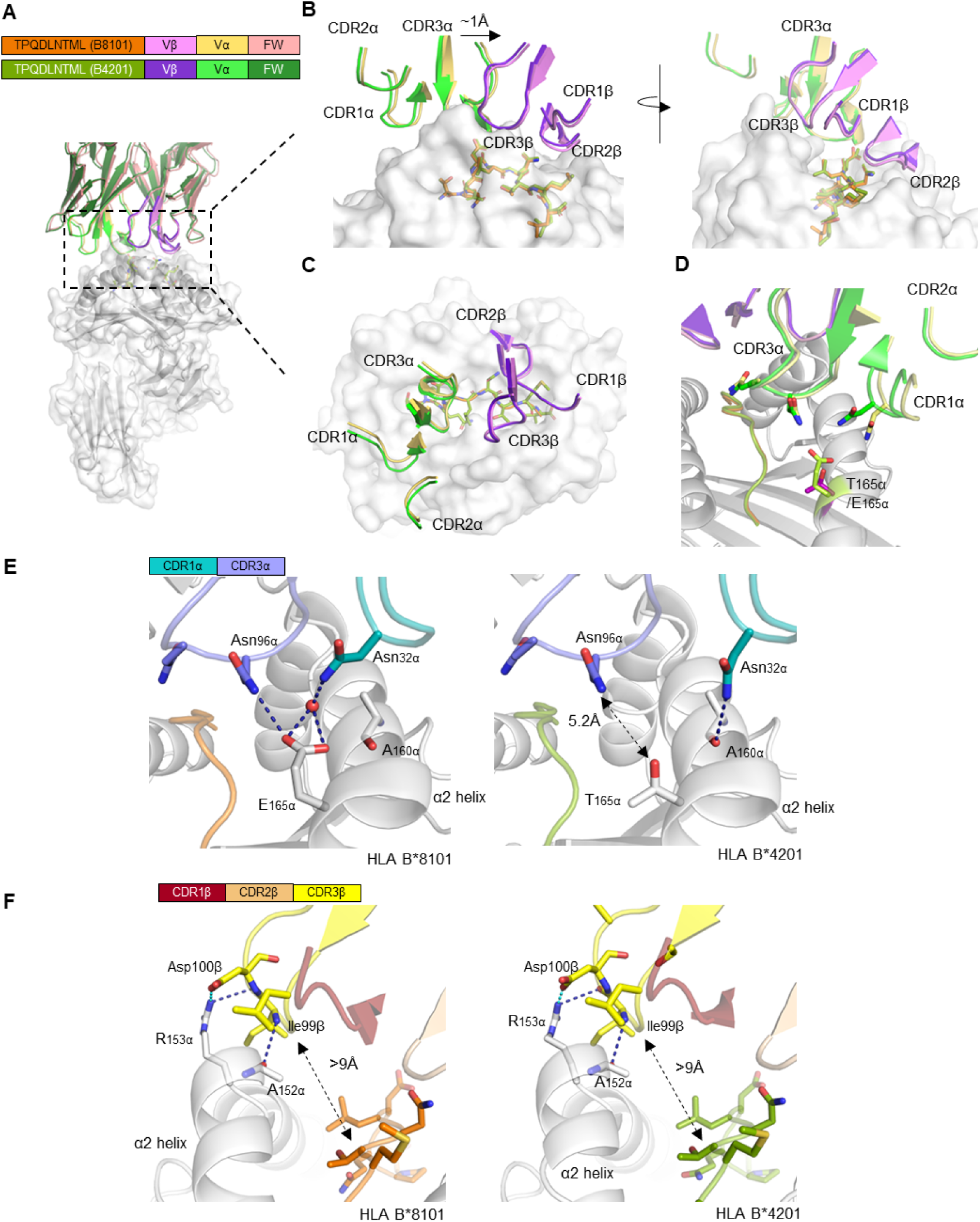
T18A adopts similar docking on B8101-restritcted TL9 and B4201-restricted TL9 but with slightly TCR twisting due to MHC polymorphism. (A). The overall view of overlapping of two ternary structures. (B) When the pHLA domains in two structures are superimposed, the CDR loops only differ by an RMSD of 0.32 Å. In the B8101 complex, the T18A TCR is positioned 1 Å closer to the peptide C terminus. CDR loops of TCR are represented in cartoon, and peptides are shown in stick. (C) The top view of CDR loops of T18A above B8101 and B42 01 molecules is shown. (D) Overlap of the polymorphic region E165α/T165α and the nearby CDR3α and CDR1α loops. (E) MHC polymorphism results in different interactions with TCR CDR3α and CDR1α loops and contributes to the swinging away of CDR1α in B4201 background. (F) CDR3β loops forms 1 salt bridge and 2 hydrogen bonds to HLA α2 helix in both structures, and is far away from the TL9 peptide. The deep blue dashed lines represent hydrogen bonds and the cyan dashed lines represent salt bridges. The online version of this article includes the following source data for figure 4: **Figure S3.** Detailed interactions of T18A CDR loops to MHC ligands. **Figure S4.** Comparison of the electrostatically colored surface of TL9-HLA-B8101 or -B4201 in complex with T18A binding.

CDR3α Loop spanned the antigen-binding cleft and contact with peptide and MHC α2 helix (Fig. 4E). It was noteworthy to mention that due to HLA polymorphisms (Fig. S4), Asn96 of CDR3α interacted with E165 of HLA-B*8101, but the side chain T165 of B*4201 was shorter than E165 of B*8101 with a distance about 5.2 Å so this H-bond was absent in B*4201 context. It was Asn32 of CDR1α formed hydrogen bond with A160 under the restriction of B*4201, as a compensation. In B*8101 context, Asn32 of CDR1α was swinging towards the MHC α1 helix and forming H-bond with E165 via the help of water molecule. Collectively, the slightly twisting of T18A docking on pMHC surface was an adaptation to the HLA polymorphism between B8101 and B4201 and contributed to cross-restriction.

In both complexes, CDR3β loops were located above the α2 helix of the HLA protein, and were far away from the peptide side chains with the distance about 9 Å, on average (Fig. 4F, Fig.S3). The CDR3β formed salt bridges between Asp100 and R153 of the HLA molecule, while Ile99 formed hydrogen bonds with R153 and A152 of the α2 helix. CDR3β of T18A formed strong contact with the bulge of MHC α2 helix and was one of the main contributors of the TCR-pMHC interaction, but indeed with no interactions towards the peptide, which was unexpected and intriguing.

### Broken of the traditional Jα connection to Vα in the T18A TCR extend its ability to bind different MHC molecules

Another surprising finding was that the traditional Jα-Vα connection was broken in T18A at both B8101 and B4201 ternary structures. The core of the traditional TCR Vα domain consists of two beta-sheets, typical in V domains of the immunoglobulin family (Fig. 5B). Unlike common “closed” Vα cores, in T18A, the disruption of the β strand made the core of Vα domain more “open” (Figure 5A). The lower part of the Jα-Vα interaction was destroyed, and three hydrogen bonds were broken near the conserved FGXG motif, but still preserved the interaction between the upper part of the chain. Moreover, the hydrogen bond between G99-G94 and N100-R93 fixed the lower portion of the CDR3α loop which might compensate for the broken of three hydrogen bonds. Such interruptions had been observed in mouse T cell responses, such as the “closed” conformation of the Yae62 TCR’s Vα bound to MHC I and the “open” conformation when bound to MHC II. In all of the “open” structures, the upper interaction between Jα and Vα strands was intact, but they were separated at the second glycine of the FGXG motif in a similar pattern, although different TRAV sequences were used. Conformational changes in the Vα core made it possible for the same TCR to cross-recognize multiple distinct MHCs (Fig. 5C).

**Figure 5.**
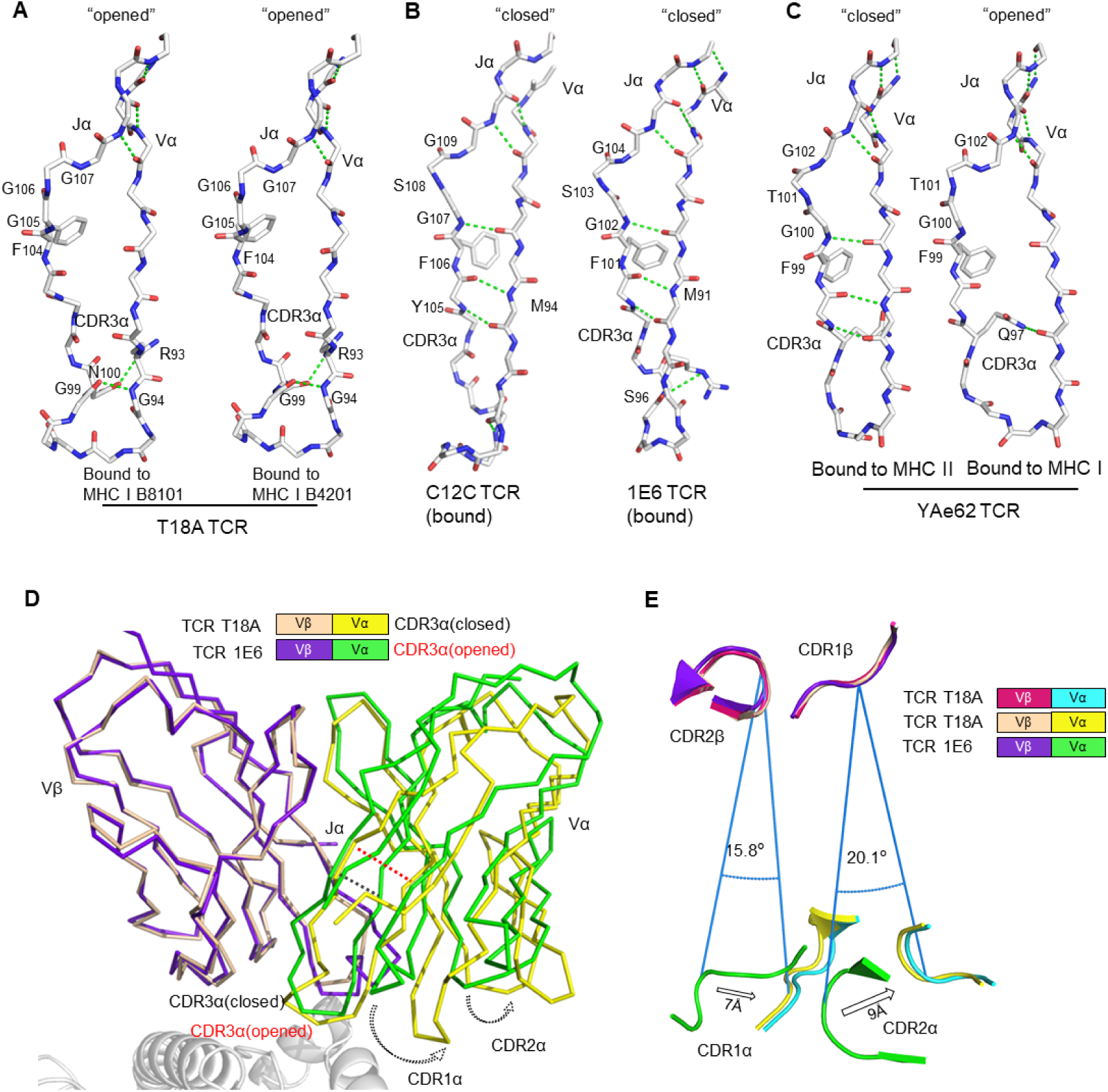
The uncommon “opened” T18A CDR3α alters the relative orientation of Vα to Vβ. (A) The “opened” conformation of the β sheet interactions between Vα and Jα of T18A when it is bound to B8101-pTL9 versus B4201-pTL9. A stick representation of the protein backbone and the side chains of the FGXG conserved motif are shown. Backbone H-bonds, as well as H-bond with R93, are shown in green. (B) The “closed” conformation of Vα-Jα interactions of C12C(Ladell et al., 2013) and 1E6(Coles et al., 2020) TCR, representing traditional CDR3α conformation in most of TCR-pMHC profiles. (C) The disruption of Vα-Jα H bonds of YAe62(Yin et al., 2011) when it is bound to MHC II versus MHC I, indicating the alteration of CDR3α could expand the ability of the TCR to adapt Different MHC Ligands. (D) The Vα and Vβ domains of T18A and 1E6 TCR are overlaid by Vβ as similar TRBV gene is used. (E) A view looking down through the TCR is shown. Relative position of CDRα loops to CDRβ loops are changed due to “opened” or “closed” CDR3α. The relative distance and angle of movement is indicated.

The direct consequence of this conformational change was to enlarge the distance between Jα and Vα, which finally led to the perturbation of the Vα domain including CDR1 and CDR2 loops, which swang away from the Vβ domain (Fig. 5D). We superimposed T18A (TRAV26-1/ TRBV12-3) and 1E6 TCR (TRAV12-3/TRBV12-4) to compare the effect of “opened” or “closed” Jα-Vα interactions on the entire TCR configuration. When Vβ domains were overlapped, the breaking of the hydrogen bond between Jα and Vα mainly affected the relative position of the Vα domain to Vβ, causing the Vα CDR1 and CDR2 rings to rotate by 15-20° relative to Vβ (Fig. 5E). The opening or closing of Jα-Vα strands above the CDR3 loop altered the relative positions of Vα and Vβ CDR1 and CDR2 loops for more than 7 Å -9 Å.

The “opened” conformation of the T18A TCR Vα core region supported a hypothesis proposed by Yin et al(Dai et al., 2008; Yin et al., 2011). They suggested that a given TCR can switch between at least three alternative conformations, depending on whether VαJα or VβJβ connections were disrupted. This hypothesis highlighted another kind of TCR plasticity, which was differed from traditional CDR residue rotamers or conformational alteration on CDR loop backbones. As Yin et al. proposed their ideas based on a cross-restriction event of mouse YAe62 TCR with MHC class I and MHC class II, the disruptions of Jα-Vα connection in cross-reactive T18A provided first clinical evidence in human natural antiviral response. This also suggested that switching between alternative conformations may be partially responsible for the alloreactivity of TCRs. Collectively, the “open” or “closed” state of Vα or Vβ core region allowed TCR to maintain the conserved interaction with MHC meanwhile expand the TCR repertoire, by enlarging or minimizing the relative distance between the Vα and Vβ domains, to adapt the different MHC components with various helical spacing and sequence.

### Unusual TCR CDR3β docking on MHC component underpins tolerance to peptide diversity

Generally, T cell receptors display high diversities in CDR3 regions to contact varied antigen peptides while less diversified CDR1 and CDR2 loops mainly contact the less varied MHC molecules. In the docking of T18A TCR toward B8101 or B4201, however, CDR3β formed few contacts to the peptide but focused on the α2 helix of MHC. This rare docking pattern was different from other cases (Fig. 6A-F, Fig. S2). Firstly, in other TCR-p-MHC complexes, CDR3β directly contacted the peptide, but such interactions were seldom observed in T18A-TL9-B8101 and T18A-TL9-B4201 complexes. Secondly, the swinging away of CDR3β let the CDR2β interact with the C-terminal of peptide, and the CDR2β, CDR3α mediated interaction with peptide was suggested to be less variable than classical CDR3β, CDR3α mediated interactions, which will constraint the conformation of the peptide. Collectively, the unusual CDR3β docking of T18A on MHC component, but not the peptide, enabled the CDR3β to avoid the distinct conformation of TL9 peptide presented by B8101 and B4201 allele, which might contribute to the cross-reactive property of T18A.

**Figure 6.**
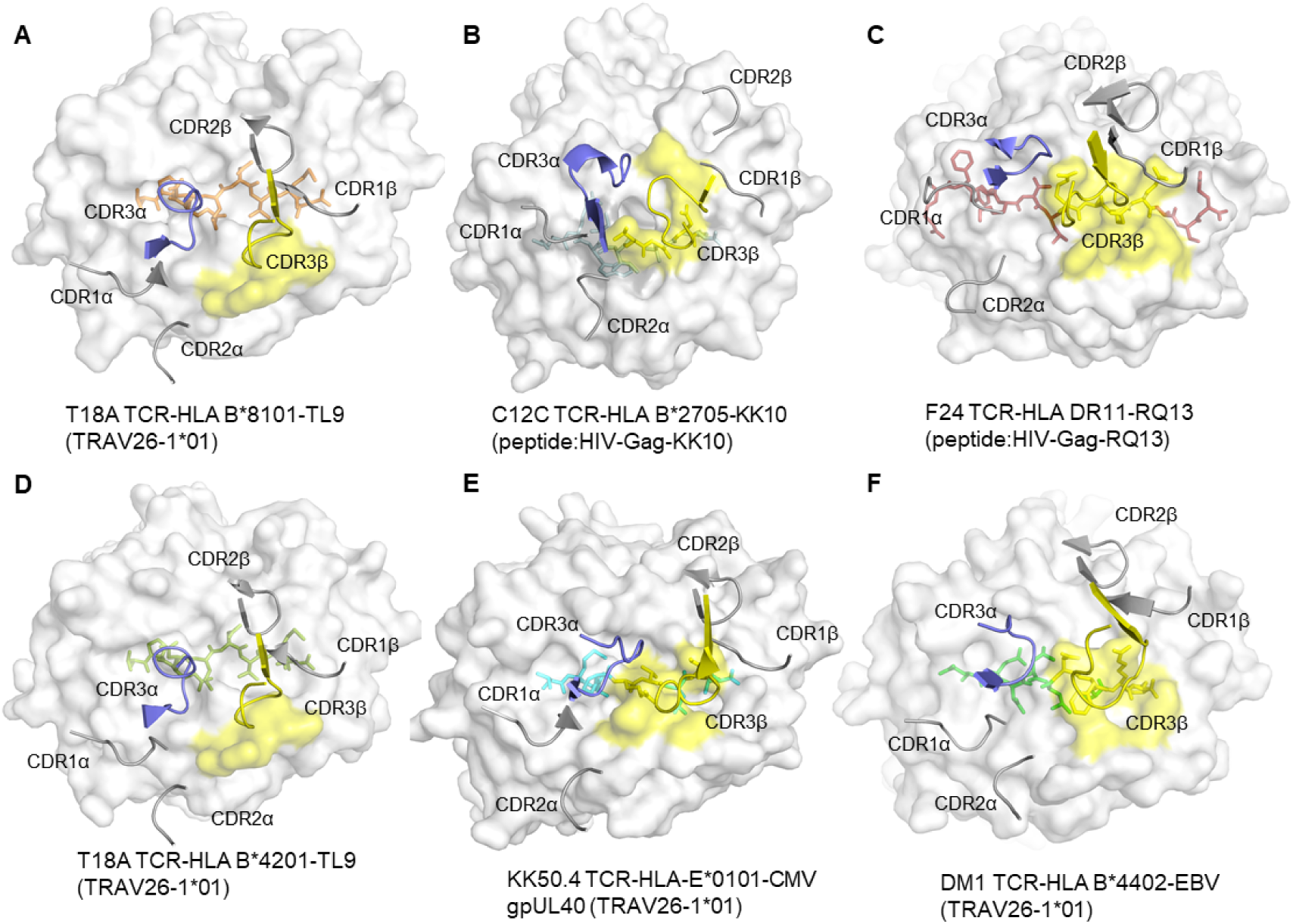
The rare docking mode of T18A CDR3β on α2 helix of the MHC but not the peptide. (A-F). The foot print of TCR CDR3β on pMHC complexes are colored in yellow from 6 different recognition profiles. Panels b-c represent the molecular mechanism of two TCR C12C and F24(Galperin et al., 2018) which are also involved in HIV immune responses; and e-f represent TCR KK50.4(Hoare et al., 2006) and DM1(Archbold et al., 2009) using similar TRAV segment as T18A TCR. Unlike other 4 structures, CDR3β of T18A recognize MHC residues instead of peptide residues; and the C terminal of TL9 is recognized by CDR2β loops. Comparison to 129 TCR-p-MHC PDB profiles confirm that the CDR3β docking mode of T18A is unique. Peptide in each panel is shown in stick, CDR loops are shown in cartoon, and MHCs are shown in surface view. The online version of this article includes the following source data for figure 6: **Figure S2.** Comparison of TCR docking between T18A and C12C reveals the leaning towards HLA α2 helix of T18A TCR.

We examined reported TCR-pMHC ternary structures from IEDB/3Dstructure database(Ehrenmann et al., 2009; Ehrenmann and Lefranc, 2011; Kaas et al., 2004; Lefranc et al., 2009) and PDB database(Burley et al., 2021), to obtain whether this CDR3β docking may have been present but not analyzed in the published data. We checked more than 260 published mouse and human TCR structures, involving 129 different TCRs (Table S4). In all of these, CDR3β interacts with peptide and MHC ligands, most mainly focused on the peptide. However, CDR3β of T18A was unique, which was far away from the peptide but formed rigidly interaction with MHC ligand. This remarkably rare characteristic of T18A extended its tolerance to mutated peptides and might be related to the delayed viral escape in the clinic.

The ‘opened’ or ‘closed’ state of the Jα-Vα or Jβ-Vβ connection was also superimposed separately in 129 different TCRs. In most of these, the β strand near the conserved FGXG motif had the conventional ‘closed’ position, but 14 TCRs had the ‘opened’ Jα-Vα connection (PDB 1MI5, 5D2N, 6AVF, 4GG6 et.al)(Broughton et al., 2012; Chan et al., 2018; Kjer-nielsen et al., 2003; Yang et al., 2015) and only one (PDB 1KJ2)(Reiser et al., 2002) had the ‘opened’ Jβ-Vβ connection. This unexpected result was not explored in these published structures, indicating the ‘opened’ or ‘closed’ CDR3α or CDR3β loops offer alternate conformations for the TCR structures and may extend the size of repertoire of a given TCR.

### High-affinity T18A TCRs bind to TL9 or TL9 escape variants under B8101 or B4201 restriction

Functional analysis and biophysical methods were then used to explore whether escape mutations on the Gag TL9 epitope and different HLA presentations affect the affinity of T18A TCR. The binding capacity of T18A TCR to different p-MHC molecules were measured by *in vitro* surface plasmon resonance (SPR). The results showed that T18A could recognize the TL9 peptide presented by B*81:01 with a high affinity (Kd≈4.7μM), and could effectively recognize some escape variants of TL9, such as 3s-TL9 and 7s-TL9 (Fig. 7A). Similarly, T18A was able to recognize TL9 peptides presented by B*42:01 with moderate affinity (Kd≈46.1μM), as well as some escape variants of TL9, further supported the dual-reactivity of TCR T18A. Through the native-PAGE assays, distinct migrations were observed of the complexes formed by T18A and B8101-TL9, B4201-TL9, B8101-3sTL9, and B8101-7SsTL9 on the gel (Fig.S5), which verified the results of the affinity measurement, that is, T18A had a high affinity against TL9 epitope of different restriction and mutations, suggesting the protective effect of this TCR in HLA-B*81:01 population.

**Figure 7.**
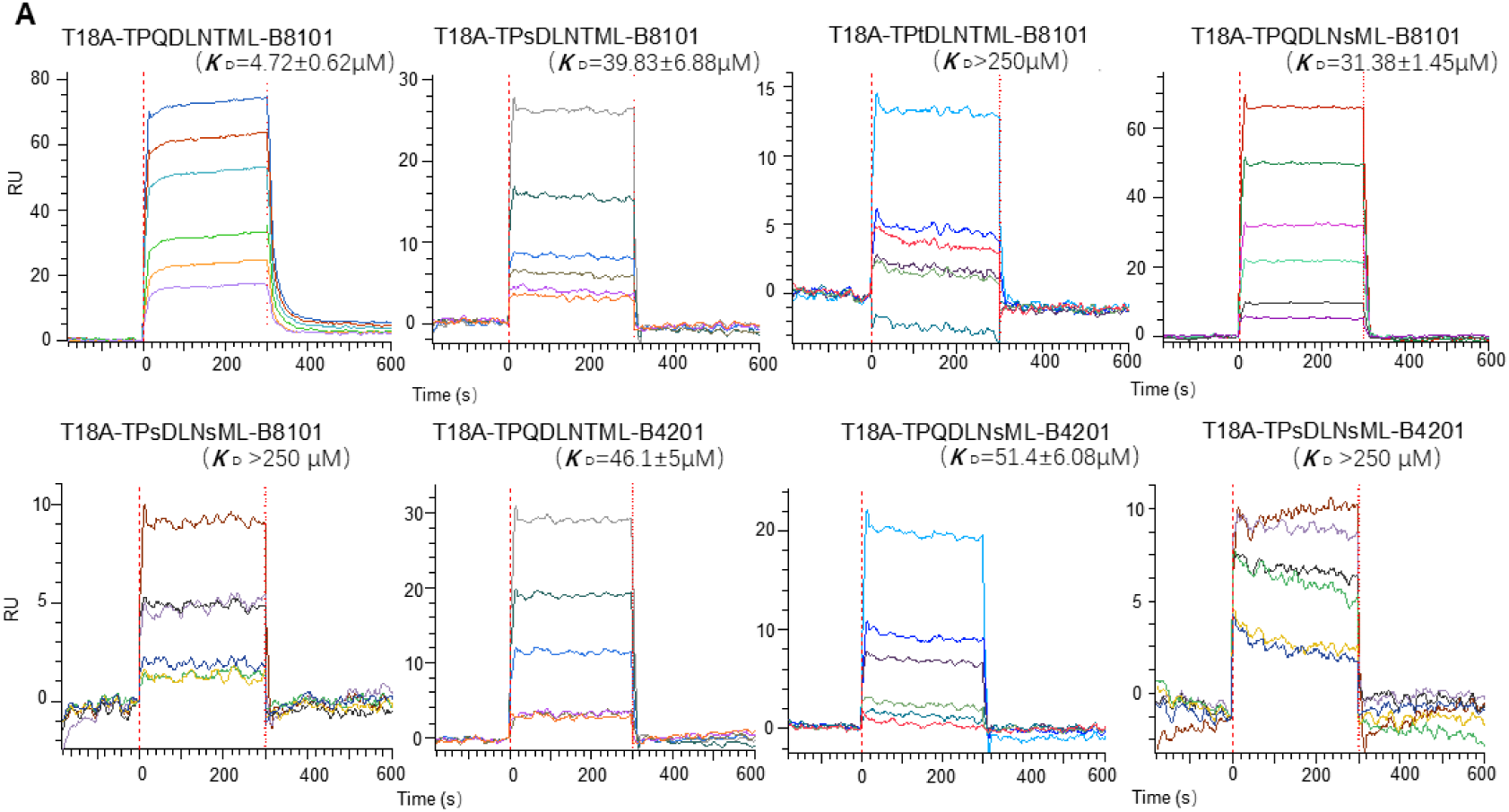
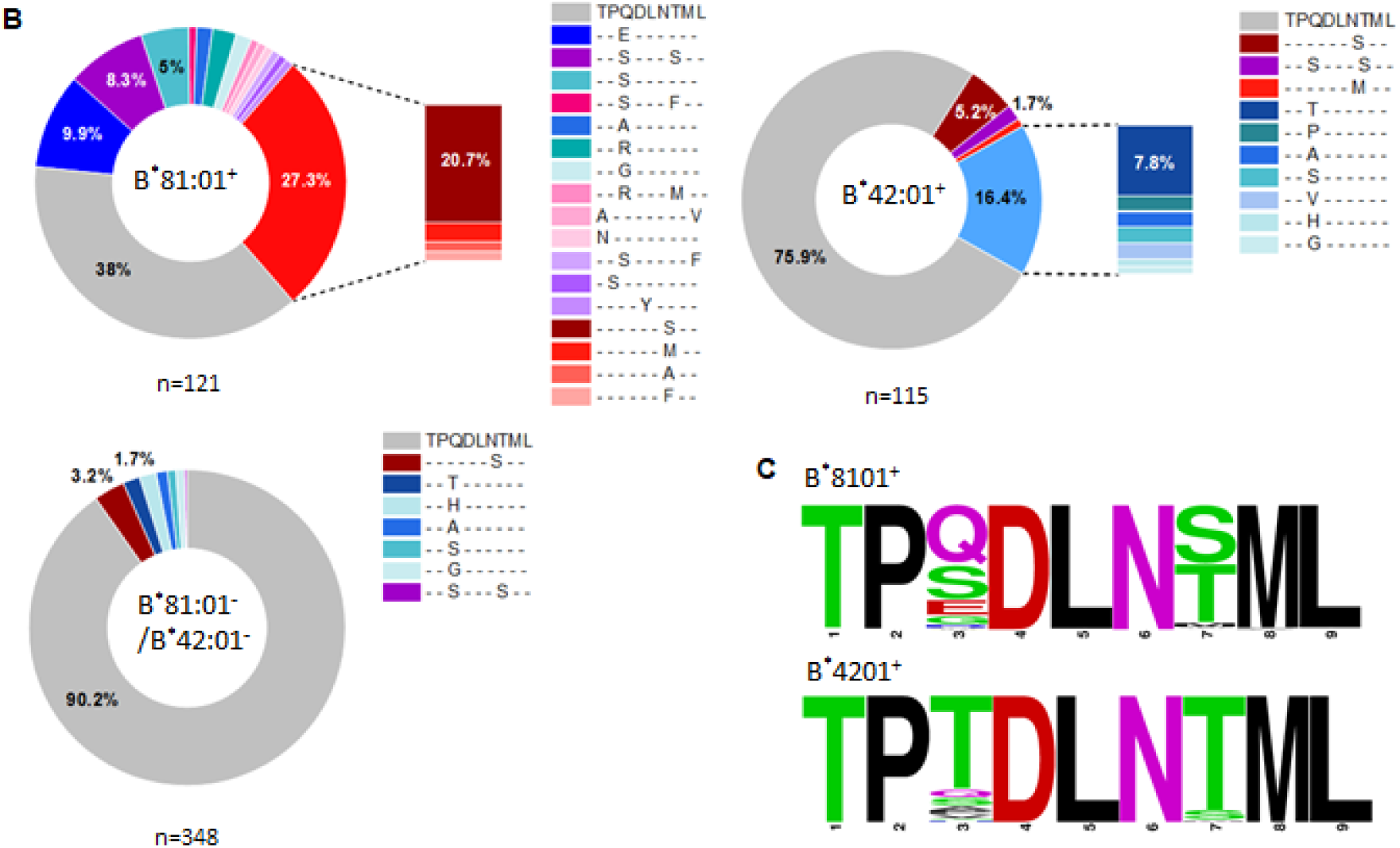
The affinity measurement by SPR and differential escape patterns in TL9 epitope under B8101 and B4201 context. (A) SPR binding data for T18A TCR recognition of the wildtype (WT) and popular mutated TL9 presented by B8101 and B4201. KD values range from 4.7 μM for the WT TL9 peptide to >250 μ M for the TPsDLNsML peptide (see also Table 1, Fig. S6). (B) HLA-associated variation in TL9-Gag in B8101-positve, B4201-positive, and B8101/B4201-negative HIV infected patients. (C) Different escape modes in TL9 epitope is illustrated as Sequence Logo, demonstrating TL9 mutation in B8101 background is located at position 3 and 7, while in B4201 background is located at position 3. The online version of this article includes the following source data for figure 7: **Figure S5.** Native-PAGE confirms the dual-reactivity of TCR T18A. **Figure S6.** Binding curves determined by SPR for mono-reactive TCRs and TL9 mutants. **Figure S7.** Mutation characteristics of Gag TL9 epitope of HIV-1 in African population. **Table S3.** HLA-associated variation in TL9-Gag from studies in last decade.

Besides, obvious differences were established for the capacity of escape mutants of TL9 epitope between mono-reactive TCR and dual-reactive TCR (Table 1, Fig.S6). Although the B*81:01-derived, mono-reactive T11A also had a strong affinity for wild-type TL9 (Kd≈4.9μM), its ability to bind mutated TL9 was weaker than that of T18A, such that only one significant binding is confirmed against TL9 mutants. On the other hand, B*42:01-derived, mono-reactive TCR T7A showed no obvious response to either wild-type or mutated TL9, as also evidenced by the native-PAGE results (Fig.S5). Since the dominant TRBV12-3*01 was used in all three TCRs mentioned above, and T18A CDR3α dominated peptide interactions but not CDR3β, we speculated that the VDJ rearrangement of the TCRα chain played a key role on the capacity of TL9 escape mutants. Finally, the direct binding assays confirmed that dual-reactive TCR T18A had functional superiority on binding the immunodominant epitope TL9, effectively adopt many escape mutants, and possibly exerting greater selection pressure than mono-reactive TCRs.

**Table 1.**
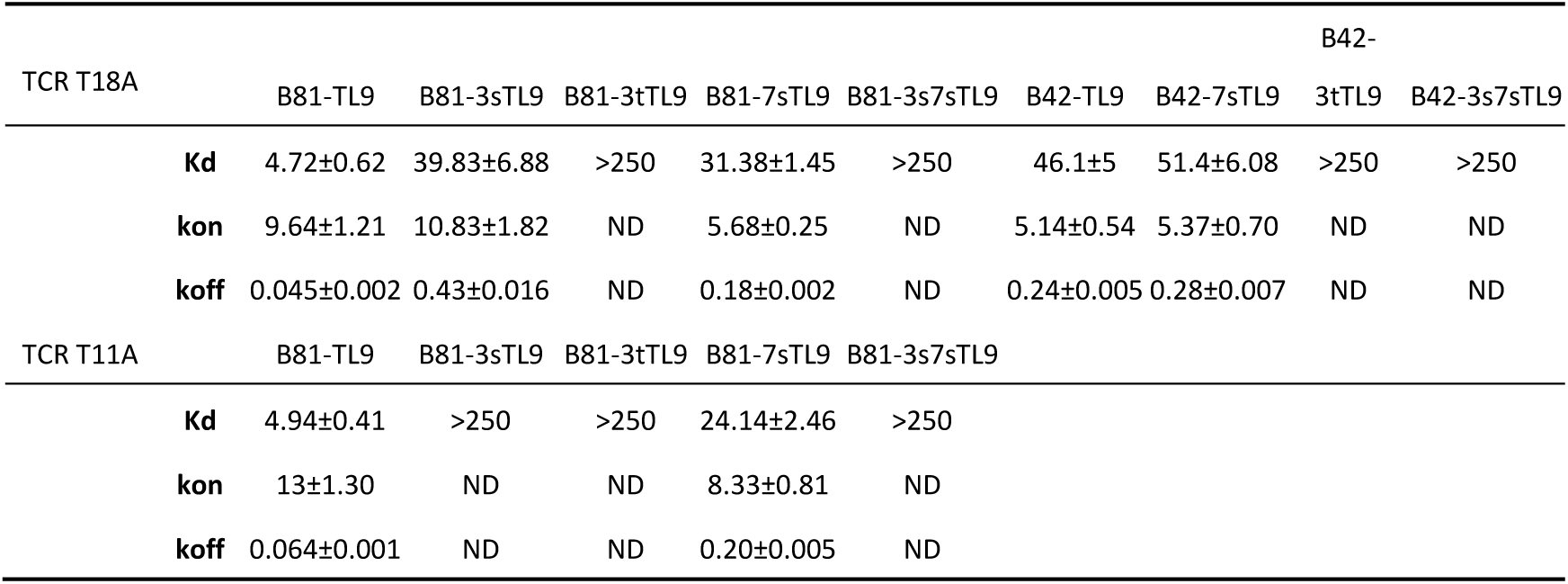
Measurement of TCR-pMHC affinity. The data of B8101-derived, dual-reactive TCR T18A (up panel) and B8101-derived, mono-reactive TCR T11A (lower panel) against HLA presented WT TL9 or mutant TL9 are listed. Kdeq is in uM; kon is in M^−1^ s^−1^ × 10^4^; koff is in s^−1^; ND, not determined; NR, no obvious responses. The error representatives the SD of triplicate experiments (n ≥ 3).

### HIV-1’s different adaptation at TL9 epitope in patients of various HLA contexts

The differences in CD8+ T cell-mediate immunity may also influence the evolution of the TL9 epitope itself. We collected the sequencing files of >3000 HIV-1 C-clade infected patients(Currier et al., 2006; Dorrell et al., 2001, 1999; Frater et al., 2007; Geldmacher et al., 2009b; Kloverpris et al., 2012; Kløverpris et al., 2016, 2015a, 2015b; Leslie et al., 2006a; Ntale et al., 2012; Payne et al., 2014) and dissected the HLA-driven differential selection pressure (Fig. 7B, Fig.S7, and table S3). The TL9 epitope of HIV-1 had a significantly higher proportion of mutations in HLA B*81:01 or HLA B*42:01 cohorts than in individuals without any of two alleles (p<0.001, t-test). It was confirmed that the TL9 epitope did run different mutation patterns across the two HLA populations.

In the context of HLA-B*81:01, the TL9 epitope mutations were mainly located at position 3 or 7 of the peptide, and the most preferred mutations were 3s-TL9 and 7s-TL9 (Fig 7C), respectively. Under the background of HLA B*42:01, the mutations in the TL9 epitope focus on position 3, and the most preferred mutation was 3t-TL9. The affinity measurement showed that mutations on these two sites of TL9 peptide could significantly reduce the affinity of TCR to pMHC molecule. Structural evidence showed that these two sites in the T18A TCR system were oriented toward the antigen-binding cleft regardless of the HLA restriction, and position 3 worked as a secondary anchored residue (Fig. 2C). This suggested that the decreased capability of T18A TCR to the mutant epitopes may mainly due to the decreased binding affinity of HLA molecule to TL9 variants. The occurrence of different HLA-specific adaptation patterns at TL9 epitope and significant differences in the affinity of TCRs indicated the qualitatively unique CTL responses induced by closely related HLA in anti-viral immunity.

## DISCUSSION

A population of dual-reactive T cells associated with lower plasma viral load following HIV-1 infection is identified by Brockman and Ndhlovu et.al(Ogunshola et al., 2018), but why these TCRs can be cross-reactive with distinct alleles and how it could help to defend chronic infections remain a mystery. Our work found essential characteristics associated with alloreactivity, which might illustrate the mechanism underpinning this biological event. Firstly, the CDR3β of T18A surprisingly focusing on recognizing the α2 helix of the HLA molecule but not the peptide, which is distinct to most known TCR recognition patterns. This unique usage of CDR3β helps the alloreactive TCR to ignore the conformational difference of the peptide, in this case, which is caused by HLA polymorphism between B8101 and B4201. Secondly, the uncommon ‘opened’ state of Jα-Vα connection changes the relative orientation of Vα to Vβ, which expands the TCR’s adaptation ability with various MHC alleles. Thirdly, the rigidity of T18A also contributes to the alloreactivity. The rigidity is performed in at least three aspects: one is the H-bond net in CDR3α makes it more relative steady than the peptide ligand; another is the usage of CDR2β instead of CDR3β increases the rigidity of TCR towards the peptide, and the third is the short CDR3 length of T18A restrict the flexibility of CDR3 loops. The rigidity of T18A TCR leads to the TL9 peptide adapts its plastic conformation to TCR docking but not in reverse.

Another intriguing question is why the dual-reactive T cells are correlated with better control against HIV-1 infection, instead of the mono-reactive T cells. This could be explained by viral escaping on the immune-dominant TL9 epitope. Our results had revealed that the TL9 peptide is flexible in the antigen-binding cleft, so besides attenuating epitope presentations, mutations on the TL9 could possibly challenge the effective TCR recognition by changing residues facing the TCRs. Considering TL9 peptide exposed distinct residues to T cells with B8101 or B4201, thus it is a great challenge for mono-reactive T cells to cope with diversified interaction surfaces. Thus the escape mutations on the TL9 epitope might sometimes change the peptide conformation and escape the pre-existing effective T cells. However, this escape strategy could be blocked by cross-reactive T cells.

From a structural perspective, the absence of CDR3β in interactions toward peptides and intensive interactions of CDR3β toward MHC make the dual-reactive TCR T18A less specific but more versatile. Polymorphic alleles B8101 and B4201 do influence the conformation of the peptide, but T18A TCR overcome this challenge. Unique CDR loop usage enables T18A to tolerate different initial conformations of the TL9 epitope, and SFR assays confirm that the affinity of dual-reactive T18A TCR for TL9-HLA, especially for mutated epitopes, was stronger than that of single-reactive TCR T11A and T7A. Besides defining antigen-specificity, the affinity of TCR to pHLA is directly correlated to the toxicity and proliferative capacity of TCR-transduced T cells, which further explains the clinical benefit of the presence of dual-reactive T cells.

Of note, the polymorphism at position 165 of MHC α2 helix (glutamic acid in B8101 but threonine in B4201) explains why the affinity of T18A against B8101-TL9 is higher than that of T18A against B4201-TL9, as H-bond is formed only between 165E and CDR3α. A stronger CD8+T immune response therefore produces greater selection pressure for HIV-1 in the B*81:01 population. Interestingly, HIV-1 sequence analysis based on >2000 individuals showed that the mutation frequency of TL9 epitope in the B*81:01 expressing individuals was significantly higher than that in the B*42:01 expressing individuals and the cohort without the above two alleles. Combined together, the differential mutation pattern of HIV-1 in different HLA contexts demonstrates how the interaction of disease-specific TCR shapes the adaptation of HIV for mutations to counterstrike the immunity.

However, there remains insufficient evidence to reveal to what extent the viral evolution is shaped by the human immune system. Reasonable speculation is that this effect is more evident in RNA viruses because the high mutation rates in virus replication provide more options for the evolution of escape variants(Singh et al., 2017). Another key point is that most of the immune-selective mutations might occur on the epitopes upon the T cell-mediated immunity. Based on the molecular arm race between CD8+ T cells and HIV-1 within the epitope TL9, the influence of host acquired immunity in genomic mutations of the virus, therefore, might be underestimated, especially for those RNA viruses that are globally prevalent, such as HIV, influenza, and SARS-CoV-2.

Collectively, our findings indicated the unique usage of CDR3β strengthens the peptide tolerance of T18A, and thus increasing the capability of TL9 escape variants. These features are consistent with the better control of viral replication and delayed viral escape in B8101 individuals. Supported by these clinical and structural evidence, the dual-reactive phenotype of CD8+ T cells might be good biomarkers for viral control and with great clinical significance for immunotherapy.

## MATERIAL AND METHODS

### Peptides

The HIV Gag p24 TL9 peptide (TPQDLNTML180-188), the escape variant Q182S, Q182T, T186S, and Q182S/T186S TL9 peptide were synthesized at > 95% purity, were synthesized at GL Biochem corporation and confirmed by high-performance liquid chromatography.

### TCR and HLA Protein expression, refolding and purification

The B*81:01/B*42:01 dual-reactive T18A TCR, mono-reactive B*42:01-restricted T7A TCR, and mono-reactive B*81:01-restricted T11A TCR were bacterially expressed as previously described(Cole et al., 2008, 2006; Hellman et al., 2016). The soluble HLA-B*81:01-TL9, HLA-B*42:01-TL9 and HLA-TL9-variants forms were also produced from bacterially expressed inclusion bodies. In brief, the α- and β-chains of TCR, the heavy chain and β2m of HLA were expressed separately as inclusion bodies in a BL21 Escherichia coli strain. The inclusion bodies were washed three times and resuspended in 8M urea, then mixed into a cold refolding buffer. For TCR refolding, 1:1 ratio of α and β chains were diluted into 50 mM Tris (pH 8.3), 2 mM EDTA, 2.5 M urea, 0.5mM oxidized glutathione, and 5mM reduced glutathione. For pMHC refolding, 1:1 ratio of HLA-B*81:01 or B*42:01 heavy chain and β2m were mixed into 100mM Tris-HCL (pH 8.3), 2mM EDTA, 400mM L-arginine-HCl, 0.5mM oxidized glutathione, and 5mM reduced glutathione. Peptides were dissolved in DMSO and injected into the refolding buffer of five molar excess folds. TCR and pMHC complexes were incubated in refolding buffer for 74 h and 48h at 4 °C, respectively. TCR and pMHC proteins were dialyzed and further purified via anion exchange chromatography (HiTrap Q HP; Mono Q; GE Healthcare) and size-exclusion (Superdex 200; GE Healthcare) as describe previously(Petersen et al., 2014; Pieper et al., 2018). The purified protein was buffer-exchanged to 10 mM Tris-HCl, pH 8.0 and concentrated to 10 mg/ml for crystallization.

### Crystallization and diffraction data collection

Protein crystals of TCR-pMHC complexes were grown at 20°C using the sitting-drop vapor diffusion technique. The T18A in complex with HLA B*81:01 and Gag TL9 peptide was crystallized in the presence of 0.2 M Potassium chloride, 0.05 M HEPES, 35% v/v Pentaerythritol propoxylate (5/4 PO/OH), pH 7.5 while the T18A in complex with HLA B*42:01 and Gag TL9 was crystallized in the buffer of 0.1 M SPG, 25 % w/v PEG 1500, pH 7.0. For cryoprotection, protein crystals were soked in 20% glycerol/80% mother liquor for 15s and frozen into liquid nitrogen. Data were collected at the BL19U1 beamline at the Shanghai Synchrotron Radiation Facility and process with HKL2000. The structures were solved by molecular replacement method using PHENIX.phaser and refined by PHENIX.refine program. Manual refinement was running in Coot. The visualization of structures was performed in PyMol and the data was deposited in the Protein Data Bank with PDB ID 7DZN, 7DZM.

### Surface plasmon resonance

The SPR assays were performed as described previously(Blevins and Baker, 2017; Kurt H. Piepenbrink, Brian E. Gloor, Kathryn M. Armstrong, 2009; Riley et al., 2018). Briefly, the protein was buffer exchanged into PBS and biotinylated for 1h at room temperature. The T18A TCR was fixed on the streptavidin-coated flow-cell surface of a SA sensor chip and the pMHC complexes were used as analyte. Injected pMHC proteins spanned concentration ranges of 0.5–250 μ M, and the equilibrium affinities were measured in 10mM HEPES, pH 7.4, 500mM NaCl, 1%BSA, and 0.02%TWEEN20 at 25°C on the Octet QKe system (ForteBio). The Kd was determined by the fitting of a single-ligand binding model.

### Analysis on sequence of HIV-1 Gag TL9 epitope from subject studies

To clearly define the HLA-B*81:01 an B*42:01-mediated differential HIV-1 epitope evolution, we collected the viral sequencing profiles from published subject studies restricted to Gag TL9 epitope since 2007 to present. More than 20 literatures were obtained. Due different scope of statistics from various studies, however, we summarized all the data and divided it into two categories: a) A total of 584 HIV-1 infected individuals with clearly identified mutant residues at TL9 epitope. b). All data was combined together, a total of 3092 HIV-1 infected persons, but with less information about the mutated residue of TL9. The data was analyzed and visualize in figure 7, table S3, and figure S7.

## ACKNOWLEDGMENTS

We thank the staff of the Shanghai Synchrotron Radiation Facility (beamline BL19U1). We sincerely pay tribute to the people who have strived in the forefront of fighting against the HIV-1 pandemic and who studied this virus around the world.

## DECLARATIONS

### Funding

This work was supported by the National Natural Science Foundation of China (31870728 and 31470738).

### Competing interests

The authors declare no conflict of interest.

### Availability of data and material

The atomic coordinates and structure details reported in this work have been deposited in the Protein Data Bank, www.pdb.org (PDB ID codes 7DZM and 7DZN).

### Code availability

Not applicable.

### Author contributions

Y.L. conducted the protein expression, purification, and crystallization. D.S did the SPR assays. L.Y. and Y.L. contributed to the study design. Y.L. and D.S contributed to data analysis. Y.L. wrote the manuscript and all authors contributed to revisions.

### Ethics approval

Not applicable. No patients are involved in this study. The clinical data are cited and summarized from published papers.

### Consent to participate

Not applicable.

### Consent for publication

All authors agree with the submission of this manuscript, and the material is original research and has not been previously reported and is not under consideration for publication elsewhere.

## Supplementary Materials

**Figure S1.** Comparison of free or TCR-bound TL9 peptide when presented by HLA-B*81:01.

**Figure S2.** The bias location of T18A TCR towards MHC α2 helix. This position of TCR drives the CDR3β loop swing away from the axis of antigen-binding cleft, and leaves less flexible CDR2β to contact with peptide C-terminus.

**Figure S3.** Comparison of CDR loops interactions between the cross-restriction structures.

**Figure S4.** The APBS electrostatics is shown by a surface view, and the polymorphic residue E165 for B*81:01 versus T165 for B*42:01 are highlighted.

**Figure S5.** Native-PAGE validates that the *E.coli*-expressed T18A TCR can cross-reactive to both B*81:01-presented TL9 and B*42:01-presented TL9. These assays also confirm WT and Q3S, Q3T and T7S TL9 cross-reactivity of the T18A.

**Figure S6.** TCR binding curves determined by SPR across different HLA-B molecules presenting WT and mutated TL9 epitope.

**Figure S7.** Analysis on HIV TL9 variation in African population with different HLA alleles from published literatures.

**Table S1.** Data collection and refinement statistics of TCR-peptide–HLA-B structures.

**Table S2.** The table provides a detailed account of contact residues between the T18A TCR and B*8101-TL9 and B*4201-TL9. It is important to understand the atomic basis of the HLA-TCR interaction.

**Table S3.** Different evolution patterns of CTL specific-TL9 epitope restricted by various HLA alleles.

---

**Figure S1.**
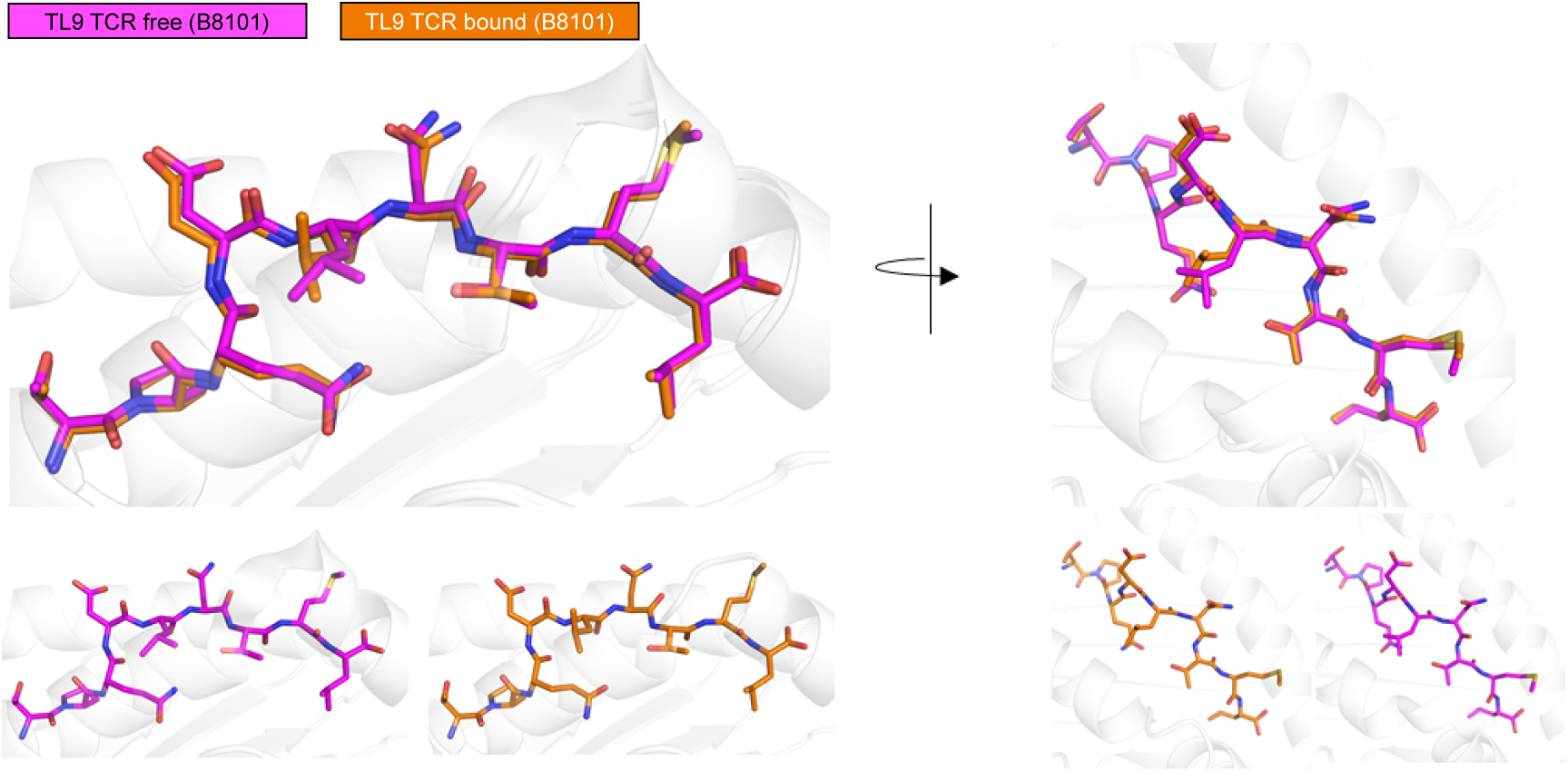
T18A engagement does not change the conformation of TL9 peptide restricted by B8101. (a) The structure of TL9 before TCR involvement is colored in magenta; The conformation of TL9 after TCR binding is colored in orange.

**Figure S2.**
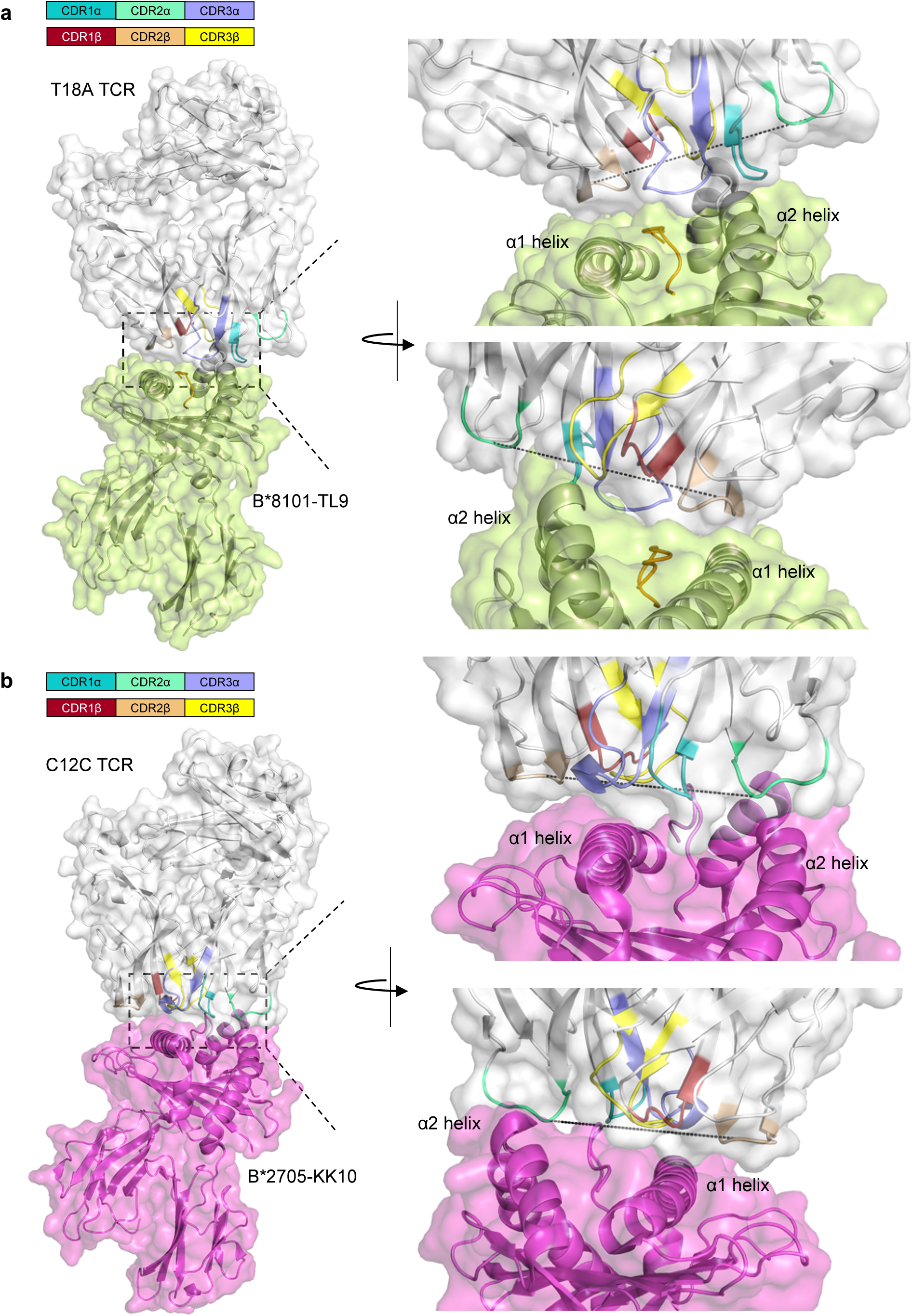
Comparison of TCR docking between T18A and C12C reveals the leaning towards HLA α2 helix of T18A TCR. (a). The surface view of T18A-B*8101-TL9 recognition. T18A positions towards to the side of HLA α2 helix, and leaves CDR3α and CDR2β to interact with peptide TL9.(b) The surface view of C12C-B*2705-KK10 recognition. KK10 is a HIV p24 Gag derived epitope, and is immunodominant in HLA-B*2705 individuals. In this recognition, TCR locates center of the antigen-binding cleft, enables the most variable CDR3β to interact with peptide.

**Figure S3.**
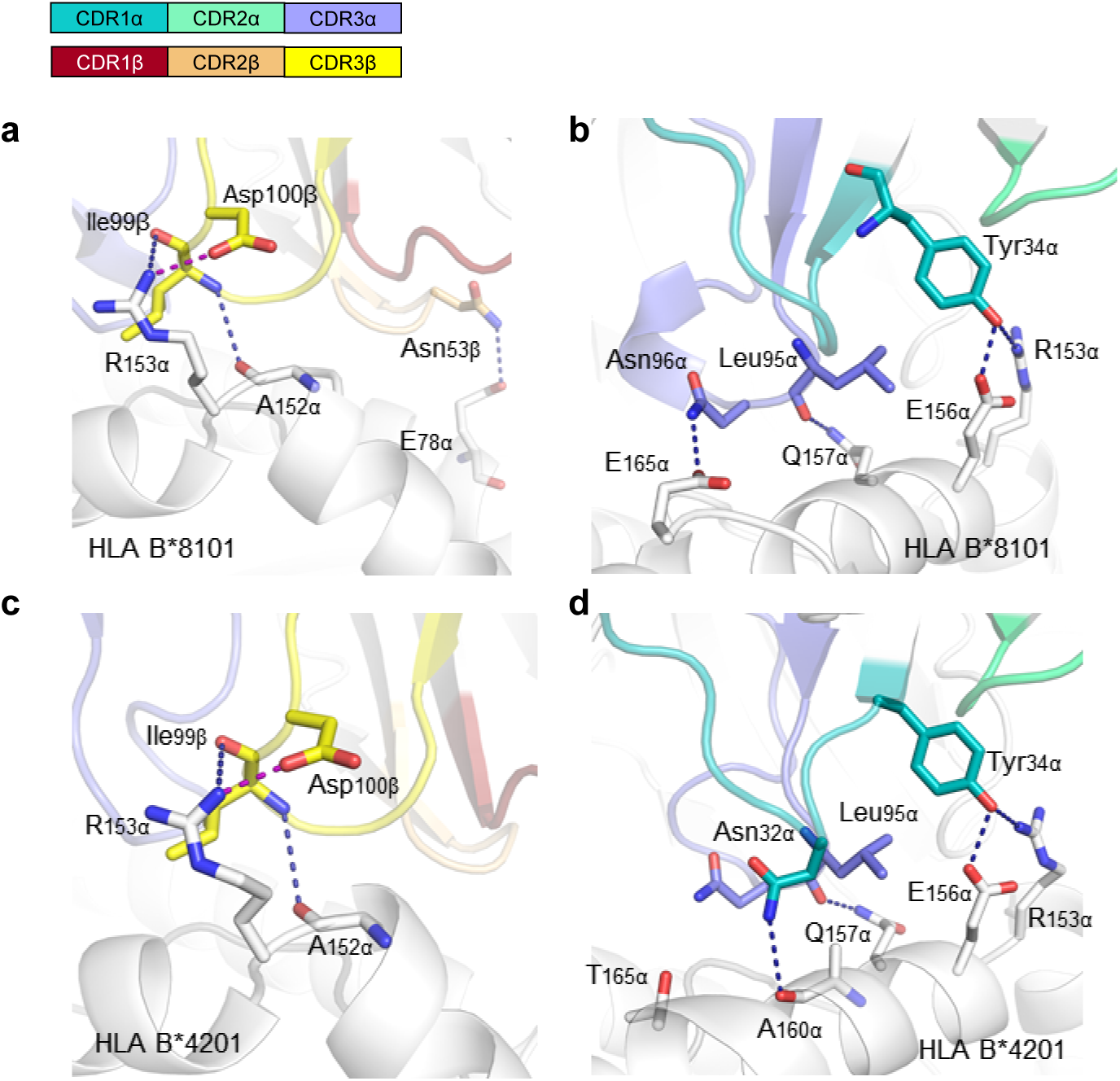
Detailed interactions of T18A CDR loops to MHC ligands. (a). Detailed view on CDR loops of the T18A β chain interact with the HLA-B8101. Hydrogen bonds and salt bridges are represented by blue or purple dashed lines, respectively. (b). T18A TCR α chain interacts with B8101 ligand. (c). CDR β loops of the T18A TCR interact with the HLA-B4201. (d). CDRα loops of the T18A TCR interact with the HLA-B4201 α2 helix.

**Figure S4.**
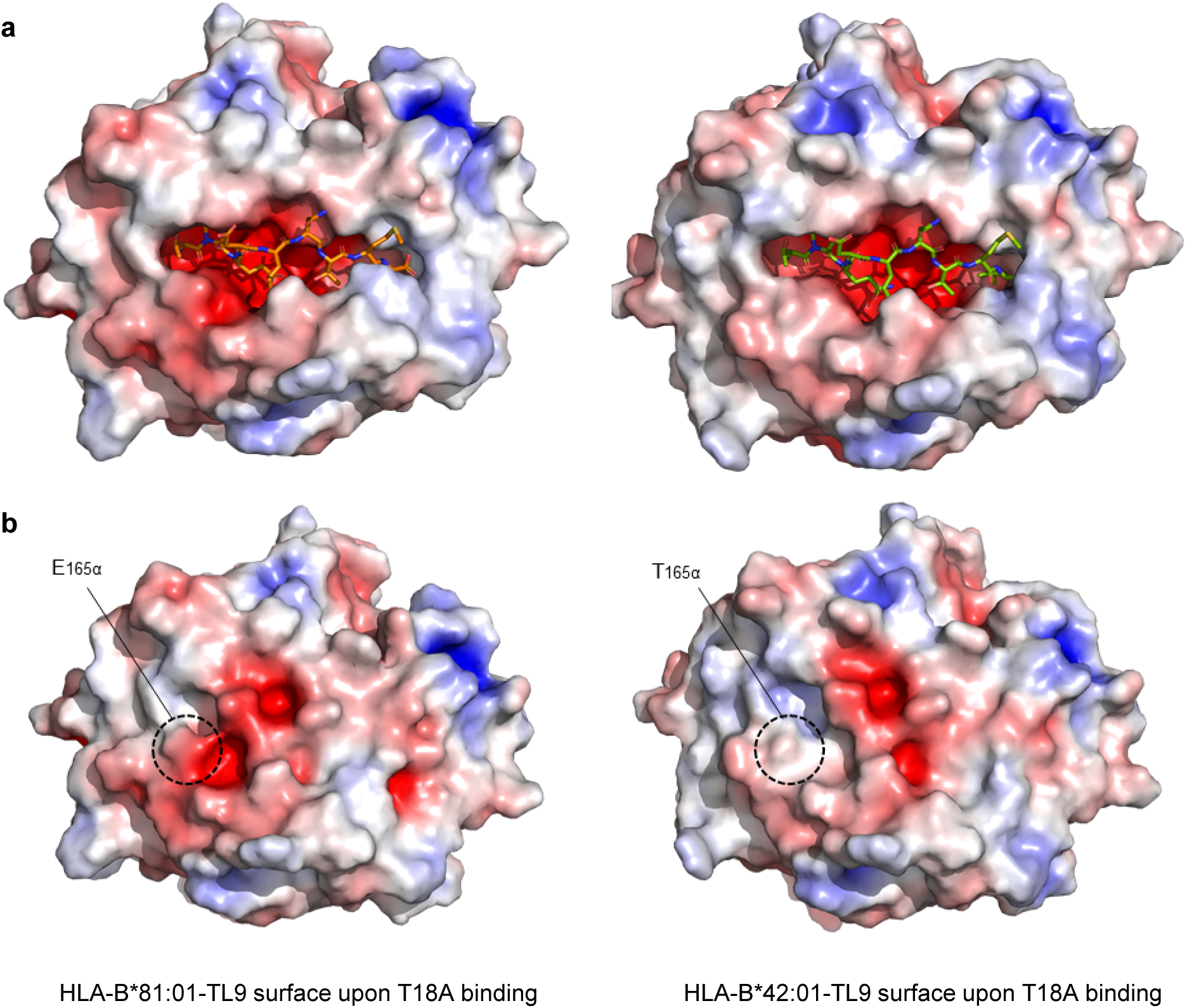
Comparison of the electrostatically colored surface of TL9-HLA-B8101 or - B4201 in complex with T18A binding. (a). TL9 in binding groove of the two structures. (b) Comparison illustrates the difference in characteristics of the TCR binding surface. The key polymorphic residue was highlighted in dashed circle. Red indicates the negatively charged and blue represents positively charged residues.

**Figure S5.**
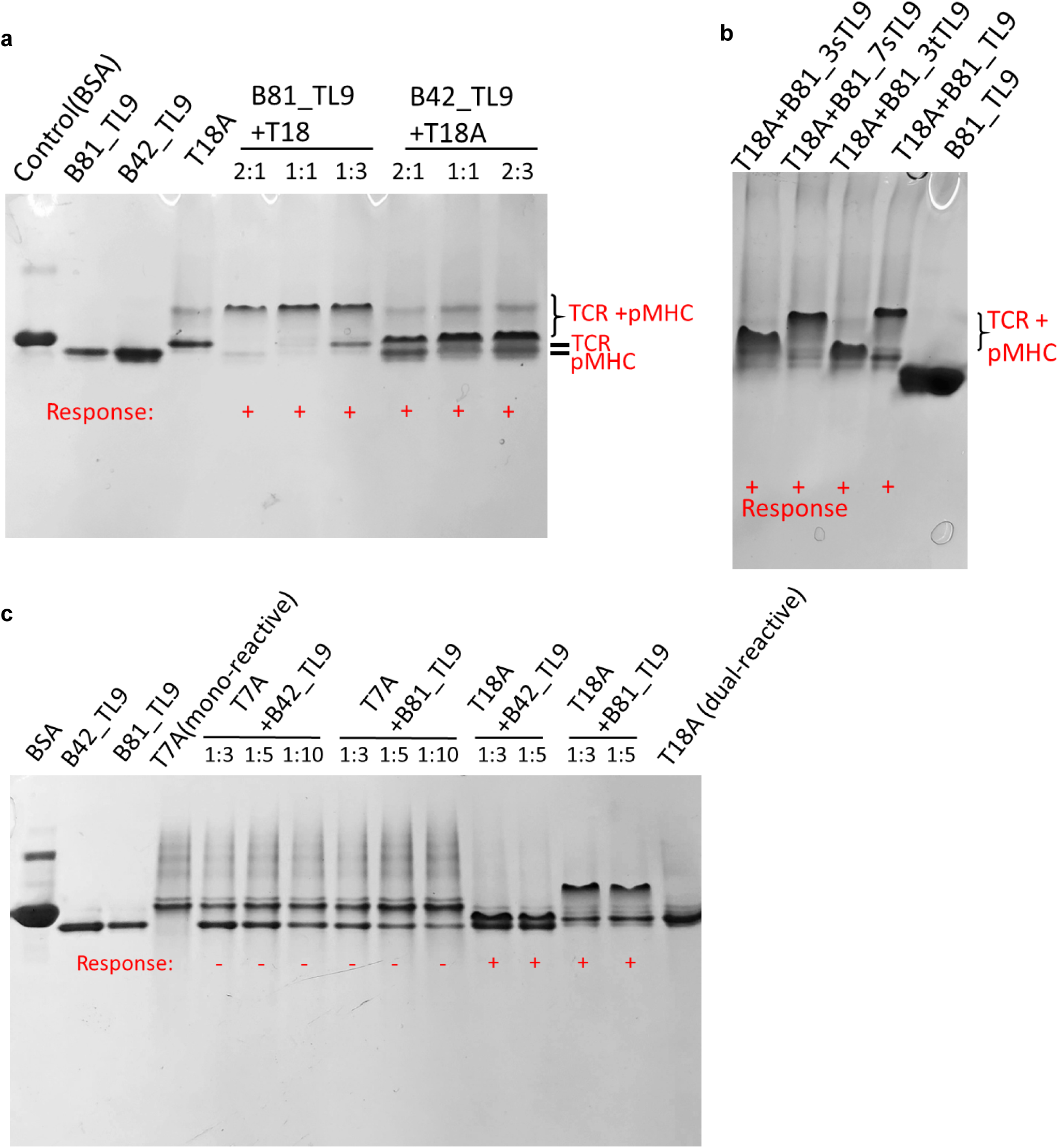
Native-PAGE confirms the dual-reactivity of TCR T18A. (a) The refolded B8101_pTL9 and B4201_pTL9 protein complexes are mixed with refolded TCR T18A separately. Compare to the print of B8101_pTL9, B4201_pTL9, and TCR only, the mixture shows positive interactions by clearly migration on the gel and demonstrate the interaction is indeed happened. This result also validate that the refolding of pMHC and TCR in vitro are successful, and with biological function to interact with each other. (b) Different migration of T18A and B81_mutated peptide confirms the ability to recognize TL9 escape variants of this TCR. (c) The negative response of single reactive TCT T7A towards B4201_pTL9 is shown, which is consistent to its weak SPR signals in affinity measurement assays.

**Figure S6.**
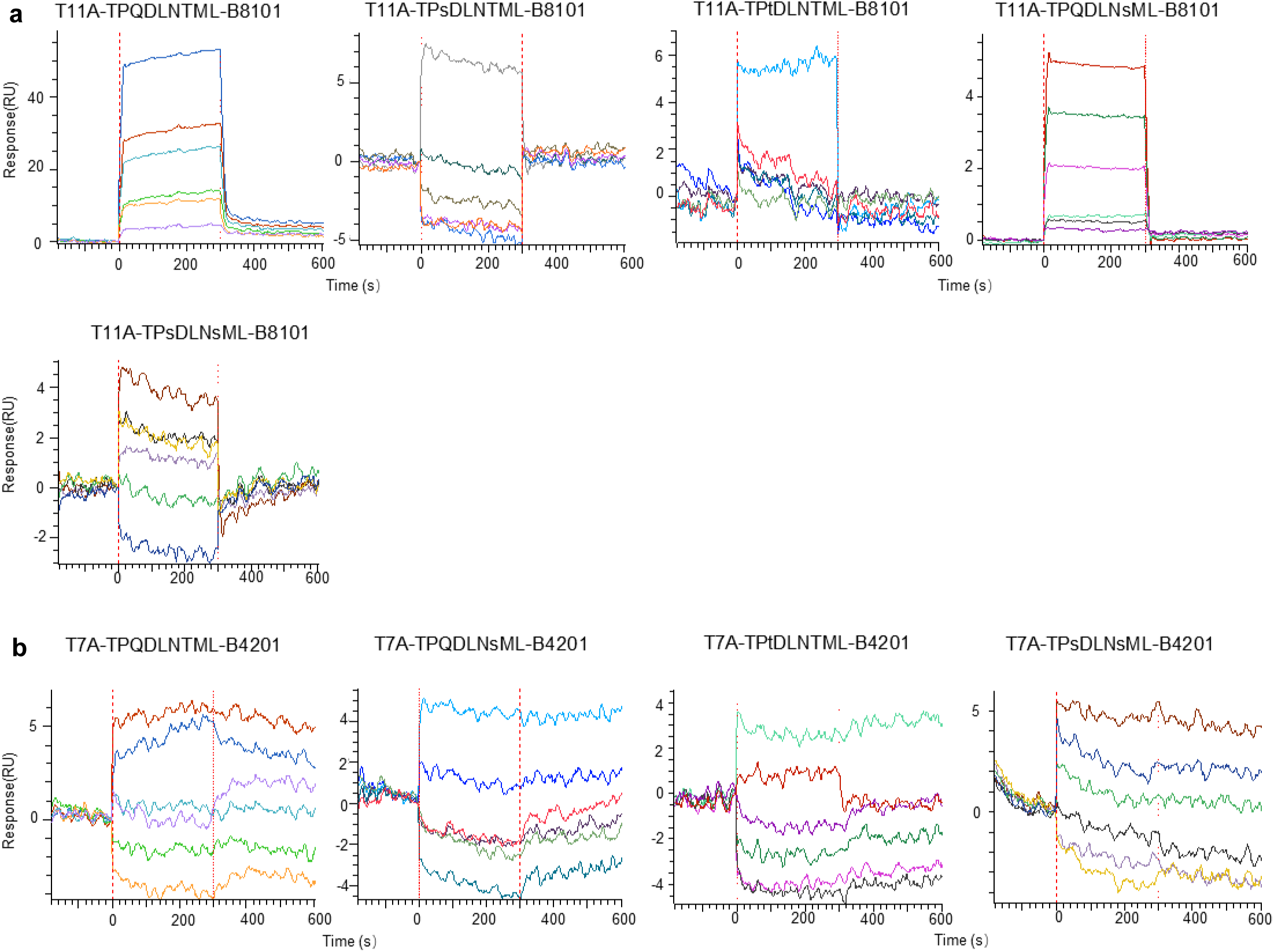
Binding curves determined by SPR for mono-reactive TCRs and TL9 mutants. The B8101-derived, mono-reactive TCR T11A (a) and B4201-derived, mono-reactive TCR T7A (b) were captured on the surface, and HLA loading WT TL9 peptide or mutated TL9 peptide were injected to the surface. The affinity was measured in response unit (RU).

**Figure S7.**
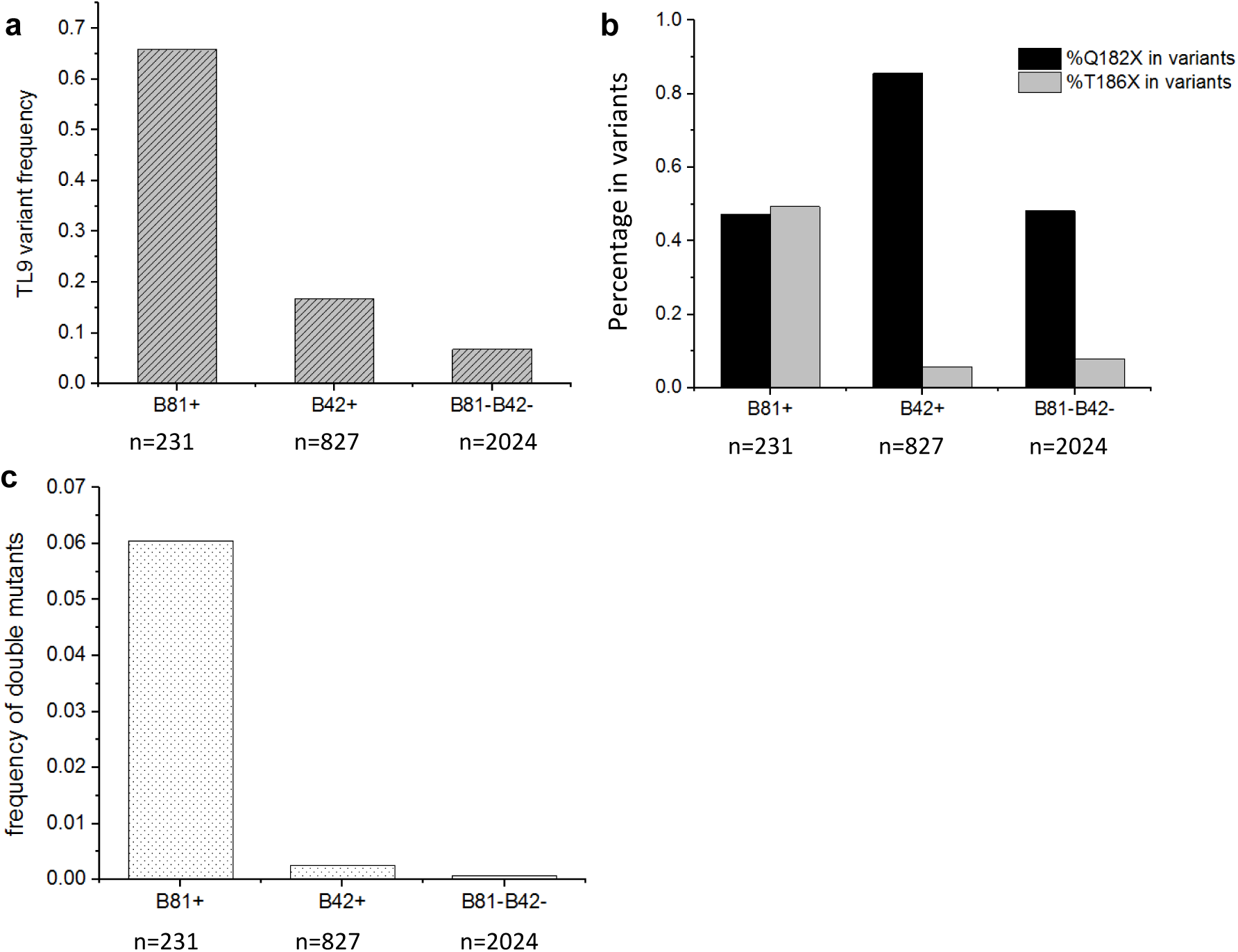
Mutation characteristics of Gag TL9 epitope of HIV-1 in African population. (a) Comparison of TL9 epitope mutation frequency between HLA B*81:01 positive, HLA B*42:01 positive, and B*81:01/B*42:01 negative population. (b) Q182X mutation proportion and T186X mutation proportion on TL9 epitope. (c) The frequency of double mutant on TL9 epitope in HIV patients.

**Table S1.** Data collection and refinement statistics of TCR-peptide-HLA complexes.

|  | T18A TCR HLA-B81-Gag-TL9 | T18A TCR HLA-B42-Gag-TL9 |
| --- | --- | --- |
| **Data collection** |  |  |
| Space group | P 43 21 2 | P 43 21 2 |
| Cell dimensions a, b, c (Å) | 93.098, 93.098, 263.051 | 96.992, 96.992, 263.839 |
| Resolution (Å) | 46.55 - 2.24 | 47.7 - 2.63 |
| Total no. of observations | 113047 (11093) | 76952 (7464) |
| No. of unique observations | 56526 (5547) | 38478 (3733) |
| Multiplicity | 2.0 (2.0) | 2.0 (2.0) |
| Data completeness (%) | 100.00 (100.00) | 100.00 (100.00) |
| I/σI | 15.84 (2.00) | 14.76 (2.25) |
| R-merge | 0.03414 (0.3732) | 0.03554 (0.2992) |
| R-meas | 0.04828 (0.5277) | 0.05027 (0.4232) |
| **Refinement** |  |  |
| Resolution (Å) | 46.55 - 2.242 (2.322 - 2.242) | 47.7 - 2.628 (2.722 - 2.628) |
| Reflections used in refinement | 56468 (5533) | 38422 (3725) |
| R-work | 0.2002 (0.2722) | 0.2176 (0.3003) |
| R-free | 0.2438 (0.3701) | 0.2661 (0.4115) |
| Number of non-hydrogen atoms | 7012 | 6802 |
| Protein | 6664 | 6716 |
| Water | 348 | 118 |
| r.m.s.d. from ideality |  |  |
| Bond lengths (Å) | 0.008 | 0.003 |
| Bond angles (°) | 0.9 | 0.64 |
| Ramachandran plot statistics |  |  |
| favored (%) | 97 | 92.7 |
| allowed (%) | 3 | 6.8 |
| outliers (%) | 0.2 | 0.3 |

**Table S2.** Contact table of T18A/HLA-B*81:01/TL9 and T18A/HLA-B*42:01/TL9

| TCR segment | TCR residues | HLA-B8101 | Type of bond |
| --- | --- | --- | --- |
| CDR1α | ASN32α | GLU165α | VDW |
| CDR1α | TYR34α | GLU156α, ARG153α | VDW, HB |
| CDR3α | LEU95α | GLN157α, GLU156α, ALA160α | VDW, HB |
| CDR3α | ASN96α | GLU165α | HB |
| CDR2β | ASN52β | THR75α, LYS148α | VDW |
| CDR2β | ASN53β | GLU78α | VDW |
| CDR2β | VAL54β | GLN74α, THR75α | VDW |
| FWβ | ILE56β | ALA71α | VDW |
| CDR3β | LEU97β | ALA151α | VDW |
| CDR3β | GLY98β | ALA152α | VDW |
| CDR3β | ILE99β | ARG153α, GLN157α, GLU156α | VDW |
| CDR3β | ASP100β | ARG153α | VDW |
| TCR segment | TCR residue | TL9 peptide | Type of bond |
| CDR3α | LEU95α | P5-LEU | VDW |
| CDR3α | ASN96α | P4-ASP | VDW, HB |
| CDR3α | ASN97α | P4-ASP, P5-LEU, P6-ASN | VDW, HB |
| CDR3α | ALA98α | P4-ASP | VDW |
| CDR2β | ASN51β | P6-ASN | VDW |
| CDR2β | ASN52β | P6-ASN, P8-MET | VDW |
| FWβ | ILE56β | P6-ASN | VDW |

| TCR segment | TCR residue | HLA-B4201 | Type of bond |
| --- | --- | --- | --- |
| CDR1α | ASN32α | ALA160α | VDW |
| CDR1α | TYR34α-OH | ARG153α-NH<sub>2</sub>, GLU156α | HB, VDW |
| CDR3α | LEU95α-O | GLN157α-NE<sub>2</sub>, GLU156α-OE<sub>1</sub>, -OE<sub>2</sub>, LEU157α, ALA160α | HB, VDW |
| CDR2β | ASN52β | GLU78α | VDW |
| CDR2β | ASN53β | GLU78α | VDW |
| CDR2β | VAL54β | GLN74α, THR75α, GLU78α | VDW |
| FWβ | ILE56β | ALA71α | VDW |
| CDR3β | LEU97β | ALA151α | VDW |
| CDR3β | GLY98β | ALA152α | VDW |
| CDR3β | ILE99β-O | ARG153α-NH<sub>1</sub>, GLU156α, GLN157α, ALA152α-O | HB, VDW |
| CDR3β | ASP100β-OD1 | ARG153α-NH<sub>1</sub> | SB, HB |
| TCR segment | TCR residue | TL9 peptide | Type of bond |
| CDR3α | ASN96α | P4-ASP | VDW |
| CDR3α | ASN97α | P4-ASP, P5-LEU, P6-ASN | VDW |
| CDR3α | ALA98α-N | P4-ASP-OD1 | HB, VDW |
| CDR2β | ASN51β-ND2 | P6-ASN-O | HB, VDW |
| CDR2β | ASN52β-ND2 | P6-ASN-O, P8-MET | HB, VDW |
| FWβ | ILE56β | P6-ASN | VDW |

VDW: Van der Waals interaction (cut-off at 4 Å), HB: hydrogen bond (cut-off at 3.5 Å), SB: salt bridge (cut-off at 5 Å).

**Table S3.**
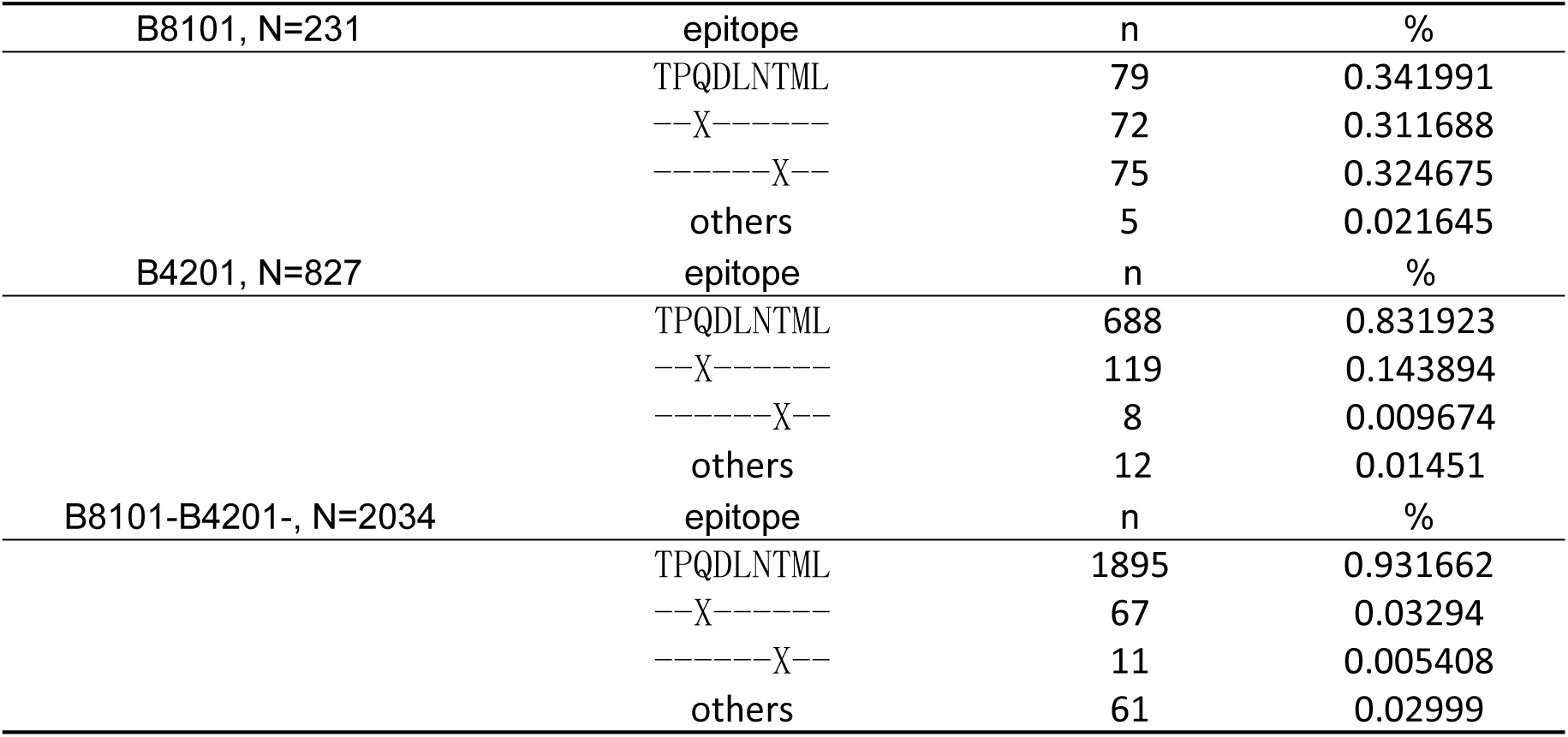
HLA-associated variation in TL9-Gag from studies in last decade.

## Notes

### Competing Interest Statement

The authors have declared no competing interest.

